# Striatal Gradient in Value-Decay Explains Regional Differences in Dopamine Patterns and Reinforcement Learning Computations

**DOI:** 10.1101/2025.01.24.631285

**Authors:** Ayaka Kato, Kenji Morita

## Abstract

Dopamine has been suggested to encode reward-prediction-error (RPE) in reinforcement learning (RL) theory, but also shown to exhibit heterogeneous patterns depending on regions and conditions: some exhibiting ramping response to predictable reward while others only responding to reward-predicting cue. It remains elusive how these heterogeneities relate to various RL algorithms proposed to be employed by animals/humans, such as RL under predictive state representation, hierarchical RL, and distributional RL. Here we demonstrate that these relationships can be coherently explained by incorporating the decay of learned values (value-decay), implementable by the decay of dopamine-dependent plastic changes in the synaptic strengths. First, we show that value-decay causes ramping RPE under certain state representations but not under others. This accounted for the observed gradual fading of dopamine ramping across repeated reward navigation, attributed to the gradual formation of predictive state representations. It also explained the cue-type and inter-trial-interval-dependent temporal patterns of dopamine. Next, we constructed a hierarchical RL model composed of two coupled systems—one with value-decay and one without. The model accounted for distinct patterns of neuronal activity in parallel striatal-dopamine circuits and their proposed roles in flexible learning and stable habit formation. Lastly, we examined two distinct algorithms of distributional RL with and without value-decay. These algorithms explained how distinct dopamine patterns across striatal regions relate to the reported differences in the strength of distributional coding. These results suggest that within-striatum differences—specifically, a medial-to-lateral gradient in value or synaptic decay—tune regional RL computations by generating distinct patterns of dopamine/RPE signals.

## Introduction

In cue-reward association learning task, it was originally found that dopamine (DA) neurons show abrupt firing response to cue but not to reward after learning, resembling the temporal-difference-reward-prediction-error (TD-RPE) in reinforcement learning (RL) (Montague et al., 1996; Schultz et al., 1997). Subsequent studies found ramping DA signals towards reward, which sustained after learning, in released DA (Howe et al., 2013; Hamid et al., 2016) and also DA neuronal activity with similar properties (Kim et al., 2020; de Jong et al., 2024), while these two signals can potentially be dissociated. Ramping DA was suggested to encode TD-RPE (Kim et al., 2020) or state value (Hamid et al., 2016; de Jong et al., 2024).

Theoretically, TD-RPE can entail sustained/ramping patterns in multiple ways (Gershman, 2014; Morita and Kato, 2014; Lloyd and Dayan, 2015; Kato and Morita, 2016; Kim et al., 2020; Mikhael et al., 2022), including the decay of learned values (hereafter referred to as value-decay), or biologically, the decay of DA-dependent plastic changes in the corticostriatal synaptic strengths (Morita and Kato, 2014; Kato and Morita, 2016) that store value memories (Reynolds et al., 2001; Samejima et al., 2005; Yagishita et al., 2014). A recent model exhibiting ramping TD-RPE (Farrell et al., 2022) also in effect assumed value-decay. Notably, according to the formulae derived for RL with value-decay (Morita and Kato, 2014), ramping TD-RPE was proportional to state value except for cue response, and so the TD-RPE and state-value accounts of ramping DA could be reconciled given the presence of value-decay.

While roles of decay/forgetting of memories have been actively studied (Davis and Zhong, 2017; Richards and Frankland, 2017; Li and van Rossum, 2020; Ryan and Frankland, 2022), value-decay is not assumed in standard RL (Sutton and Barto, 2018), but was shown to be potentially beneficial (Brea et al., 2014; Kato and Morita, 2016; Moens and Zénon, 2019; Jiang et al., 2024) and also potentially better fit behavior (Ito and Doya, 2009; Niv et al., 2015). Moreover, recent studies (Mikhael and Bogacz, 2016; Pinto and Uchida, 2023; Lowet et al., 2024) suggest that the striatum-DA system implements distributional RL (Dabney et al., 2020; Lowet et al., 2020; Bellemare et al., 2023; Muller et al., 2024), and value-decay was assumed to ensure stability against value divergence in some (Mikhael and Bogacz, 2016; Pinto and Uchida, 2023), but not other (Dabney et al., 2020; Lowet et al., 2024) distributional RL models. Value-decay could also be useful for canceling out uncertainty-induced bias (Mikhael et al., 2022) if correctly tuned (Morita and Kato, 2022).

Recent studies revealed mechanisms for forgetting/decay of memories (Shuai et al., 2010; Berry et al., 2012; Hayashi-Takagi et al., 2015; Davis and Zhong, 2017; Berry et al., 2024), including those depending on DA, in Drosophila (Berry et al., 2012; Cervantes-Sandoval et al., 2016) and mammals (Castillo Díaz et al., 2019; Gallo et al., 2022), but we could not find demonstration of the decay of value-storing cortico-striatal synaptic strengths, except for a suggestion of possible occurrence of DA-dependent forgetting in the nucleus accumbens (NAc) (Tu et al., 2019). However, there were findings that appear to imply value-decay. Specifically, monkey caudate-head (CDh) neurons rapidly developed value-encoding response within a session but lost it at the beginning of sessions in subsequent days, while caudate-tail (CDt) neurons gradually developed value response across days (Figure 6B,C of (Kim and Hikosaka, 2013)).

An emerging possibility, which we address here, is that presence/absence of value-decay could explain how different DA patterns in different regions/conditions serve for suggested distinct computations, including model-based(−like) RL (Sharpe et al., 2017; Langdon et al., 2018; Krausz et al., 2023) or RL using sophisticated state representations (Dayan, 1993; Momennejad et al., 2017; Russek et al., 2017; Stachenfeld et al., 2017), hierarchical RL (Haruno and Kawato, 2006; Ito and Doya, 2010; Botvinick, 2012; Frank and Badre, 2012; Keramati and Gutkin, 2013; Balleine et al., 2015), and distributional RL (Lowet et al., 2020; Bellemare et al., 2023).

## Materials and Methods

### Effects of value-decay under various state representations

We simulated the behavior of RL agent in a cue-reward association learning task. In each trial, upon seeing the cue stimulus, the agent entered state *S*_1_, and then sequentially and deterministically transitioned to state *S*_2_, …, and *S*_*n*_ at each time step. At state *S*_*n*_, the agent received a reward of a fixed size (*R*_*n*_ = 1) deterministically (for Figures 1 and 5) or of different sizes (*R*_*n*_ = 0.5 or 1.5) with equal probabilities (for Figure 6), and at the other states, the agent did not receive any reward (*R*_*i*_ = 0 (*i* ≠ *n*)). The number of states (*n*) was set to 5 in most simulations except for those for Figure 5, in which *n* was varied from 1 to 10. The number of trials was set to 200 for Figures 1 and 5, and 100 for Figure 6.

**Figure 1.**
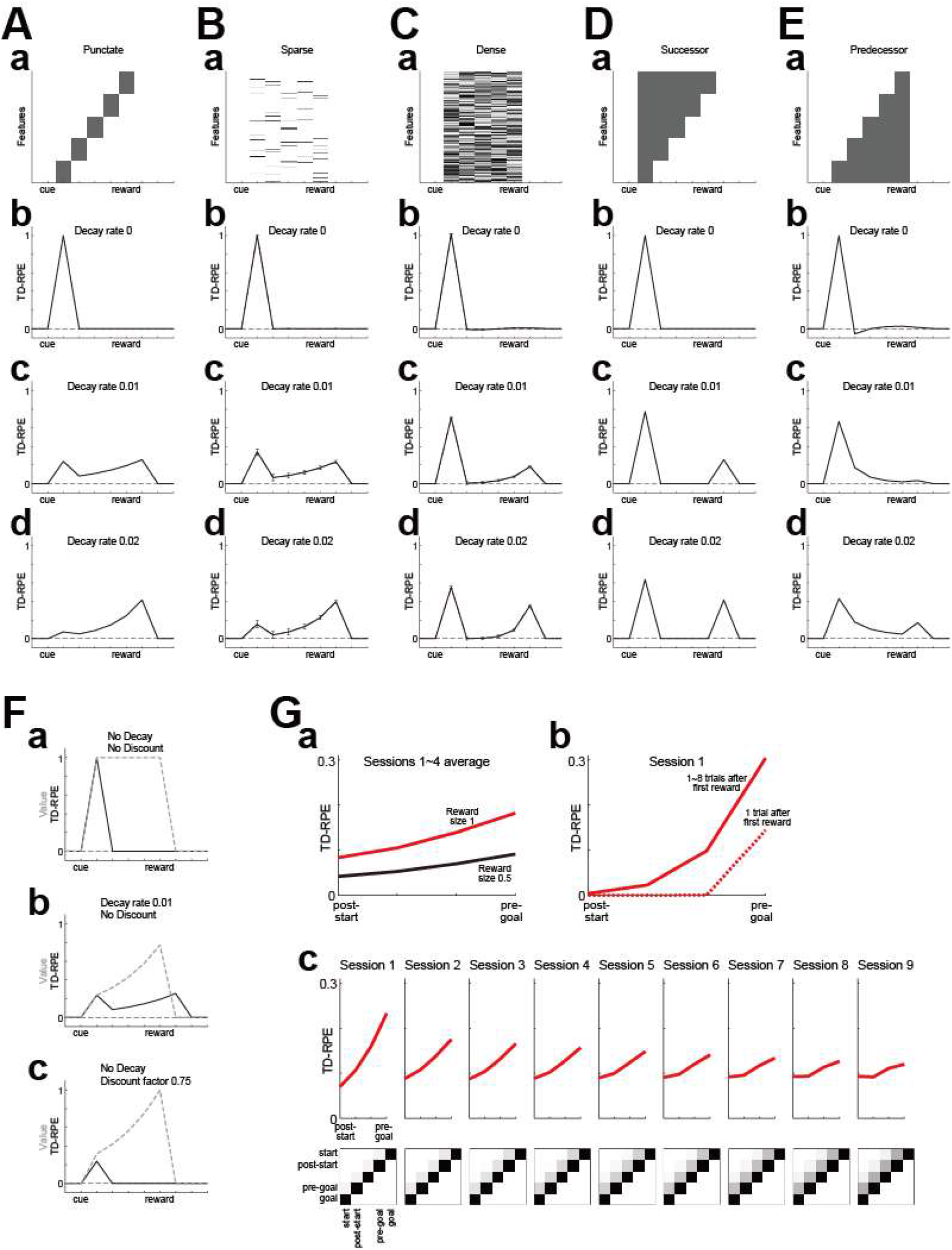
Effects of value-decay on the TD-RPE pattern under various state representations. **(A-E)** TD-RPE generated in a simulated cue-reward association learning task after learning became almost steady in cases with various state representations and rates of value(−weight)-decay. Time discount factor was set to 1 in all the cases. (A) Case where five timings from cue to reward were separately represented in a punctate manner, as schematically indicated in (a). The rate of value-decay was set to 0 (no decay) (b), 0.01 (c), or 0.02 (d). (B,C) Cases where the five timings were represented by sparse (B) or dense (C) pseudo-random feature vectors (indicated in (a)). The lines and error-bars in (b-d) indicate the mean±SD across 100 simulations. (D,E) Case where the five timings were represented by the successor representation (SR) (D) or the predecessor representation (PR) (E). **(F)** Comparison of the effects of value-decay and temporal discounting on the patterns of TD-RPE (black solid lines) and state value (gray dashed lines) in the case of the punctate state representation. **(a)** Case with no value-decay and no temporal discounting. **(b)** Case with value-decay (rate 0.01) and no temporal discounting. **(c)** Case with no value-decay and temporal discounting (time discount factor = 0.75). **(G)** TD-RPE ramping in simulated reward navigation gradually faded away when SR was gradually shaped through TD learning of SR features. **(a)** TD-RPE from post-start to pre-goal states averaged over 1st ~ 4th sessions, with each session consisting of 40 trials. The red and black lines indicate cases with reward size 1 and 0.5, respectively. Compare to DA ramps in navigation towards big and small rewards in Fig. 1e of (Guru et al., 2020). **(b)** TD-RPE averaged over 1st ~ 8th trials (red solid line) or at the 1st trial (red dotted line) after the initial trial in which reward (size 1) was obtained. Compare to DA ramps in early trials after first reward was obtained in Fig. 1j of (Guru et al., 2020). **(c)** TD-RPE (top) and state representation (feature) matrix (bottom) in 1st ~ 9th sessions. Compare with the experimentally observed fading of DA ramps over daily training in Figure 4a of (Guru et al., 2020).

We considered an agent having a learning system, which had the predicted value of each state, *V*(*S*_*i*_) (*i* = 1, …, *n*), initialized to all 0. At each time-step/state, TD-RPE:

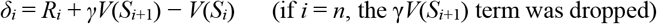

was calculated, where *γ* was the time discount factor and was set to 1 in all the simulations in Figures 1, 5, and 6 except for those shown in Figure 1Fc for which *γ* was set to 0.75 (even if *γ* was instead set to 0.9, main points were largely preserved as shown in Extended Data Figure1-1, 5-1, and 6-1). *V*(*S*_*i*_) was then updated as

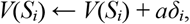

where *a* was the learning rate and was set to 0.15. At each time-step/state, the predicted values of all the states decayed (Morita and Kato, 2014) as

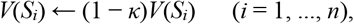

where *κ* was the decay rate and was set to 0, 0.01, or 0.02. Note that value-decay was imposed at every time-step only during the task trials.

We next considered cases where each state, *S*_*i*_ (*i* = 1, …, *n*), was represented by a 100 dimensional feature vector:

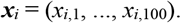

We examined cases with two different types of representation: sparse representation and dense representation. In the sparse representation, for each feature vector ***x***_*i*_, only a subset of elements (10 out of 100), which were pseudo-randomly selected, were drawn from [0 1] uniform pseudo-random numbers while the remaining elements were 0, and subsequently each feature vector was scaled so that its norm became 1. In the dense representation, all the elements of each feature vector ***x***_*i*_ were drawn from [0 1] uniform pseudo-random numbers, and subsequently each feature vector was scaled so that its norm became 1. We also considered cases where the states were represented by the successor representation (SR) (Dayan, 1993; Russek et al., 2017) or the predecessor representation (PR) (c.f., (Bailey and Mattar, 2022)). In these case, the states were represented by future or past occupancies of other (and self) states (in *n* = 5 dimension):

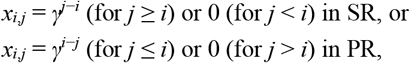

where *γ* was set to 1. In all these cases (sparse, dense, SR, and PR), predicted value of the state (i.e., state value) was calculated as an inner product of the features and a common weight vector

***w*** = (*w*_1_, …, *w*_*m*_) (*m* = 100 for the sparse and dense representations and *m* = 5 for the SR and PR) with subtraction of constants:

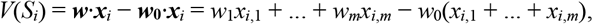

where ***w***_**0**_ = (*w*_0_, …, *w*_0_) was a constant vector, to which ***w*** was initialized, and *w*_0_ was set to 0.5. At each time-step/state, TD-RPE:

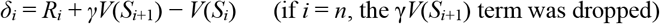

was calculated, where *γ* was set to 1 in all the simulations. The weight vector ***w*** was updated as

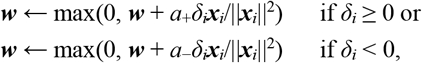

where the max operation ensured that the weights remained non-negative, and *a*_+_ and *a*_−_ were the learning rates for positive and negative TD-RPEs, respectively, which were set to 0.15 and 0.075 in Figures 1 and 5 (even if *a*_+_ and *a*_−_ were instead set to both 0.15, main points were largely preserved as shown in Extended Data Figure1-2 and 5-2) and both 0.15, 0.015, or 0.005 in Figure 6. At each time-step/state, all the weights decayed to their initial values:

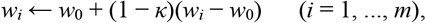

where *κ* was set to 0, 0.01, or 0.02 (note that value-decay was imposed at every time-step only during the task trials). For the cases with the sparse and dense representations, we conducted 100 simulations for each condition, and plotted the mean±SD TD-RPE (of all the simulations, or of those corresponding to each condition (reward size) for Figure 6) at each time-step/state.

We further simulated a reward navigation task in a linear track examined in a recent study (Guru et al., 2020). In fact, since we did not model sensory- and motor-related processes, our simulation of reward navigation task was just as same as our simulation of cue-reward association task described above, with the number of states (*n*) set to 5 and the reward size (*R*_5_) set to 1 or 0.5. Since the experimental study (Guru et al., 2020) reported DA response between the “start” and “goal”, which appeared to be slightly separated from the ends of the linear track (inferred from the animal’s speed in Fig. 1c of (Guru et al., 2020) and also from the schematic of the linear track in Fig. 1b of (Guru et al., 2020)), we plotted TD-RPE between *S*_2_ (named the “post-start” state) and *S*_4_ (named the “pre-goal” state). We assumed that initially each state was represented in a punctate manner but then SR was gradually shaped through TD learning of SR features. Specifically, each state *S*_*i*_ (*i* = 1, …, 5) was represented by a 5-dimensional feature vector:

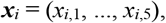

which was initialized as follows:

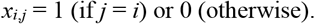

At each time-step/state *i*, TD error-vector of features:

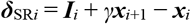

(if *i* = 5, the *γ****x***_*i*+1_ term was dropped; since ***x***_5_ was initialized to ***I***_5_, ***δ***_SR5_ was always **0**) was calculated, where ***I***_*i*_ was a 5-dimensional vector whose *i*-th was 1 and the other elements were 0 and *γ* was set to 1. The feature vector ***x***_*i*_ was updated as:

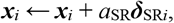

where *a*_SR_ was the learning rate for feature update and was set to 0.001. With these gradually updated state representations, calculation of state values and update of value-weight vector ***w***, including its initialization to ***w***_**0**_ = (0.5, …, 0.5) and decay, were conducted in the same manner as in the case with (constant) SR described above. *a*_+_ and *a*_−_ were set to 0.15 and 0.075, respectively, and *κ* was set to 0.01. The number of trials was set to 360, which was assumed to divided into 9 sessions with each session consisting of 40 trials.

In these simulations, inter-trial-intervals (ITIs) were not modeled, and so value-decay in ITIs was not considered. However, value-decay-induced ramping of TD-RPE can in principle occur regardless of whether value-decay occurs only during task trials or also during ITIs. In our previous studies, we assumed value-decay at every time-step in both task trials and ITIs (Fig. 9 of (Morita et al., 2013)) or value-decay once at every trial rather than at every time-step (the first part of (Morita and Kato, 2014)) and showed that TD-RPE ramping occurred in both cases. On the other hand, if value-decay is biologically implemented by activity-dependent synaptic mechanisms, value-decay during ITIs can be weak because pre-synaptic cortical activity representing the task states can be weak during such periods.

### Effects of value-weight-decay in the model that learns state representation and value

We examined the effects of value-weight-decay in the online value-RNN (oVRNN) model (Tsurumi et al., 2025) that simultaneously learns state representation and state value. Specifically, we adapted a version of oVRNN with random feedback and biological constraints (“oVRNNrf-bio” in (Tsurumi et al., 2025)). oVRNN, as well as the original value-RNN with BPTT (Hennig et al., 2023; Qian et al., 2025), consists of the observation units, an RNN, a readout unit, a reward-encoding unit, and a TD-RPE-encoding unit, which are presumed to be implemented in the sensory cortex, association/prefrontal cortex, striatum, neurons encoding reward information, and DA neurons, respectively (Fig. 2A).

**Figure 2.**
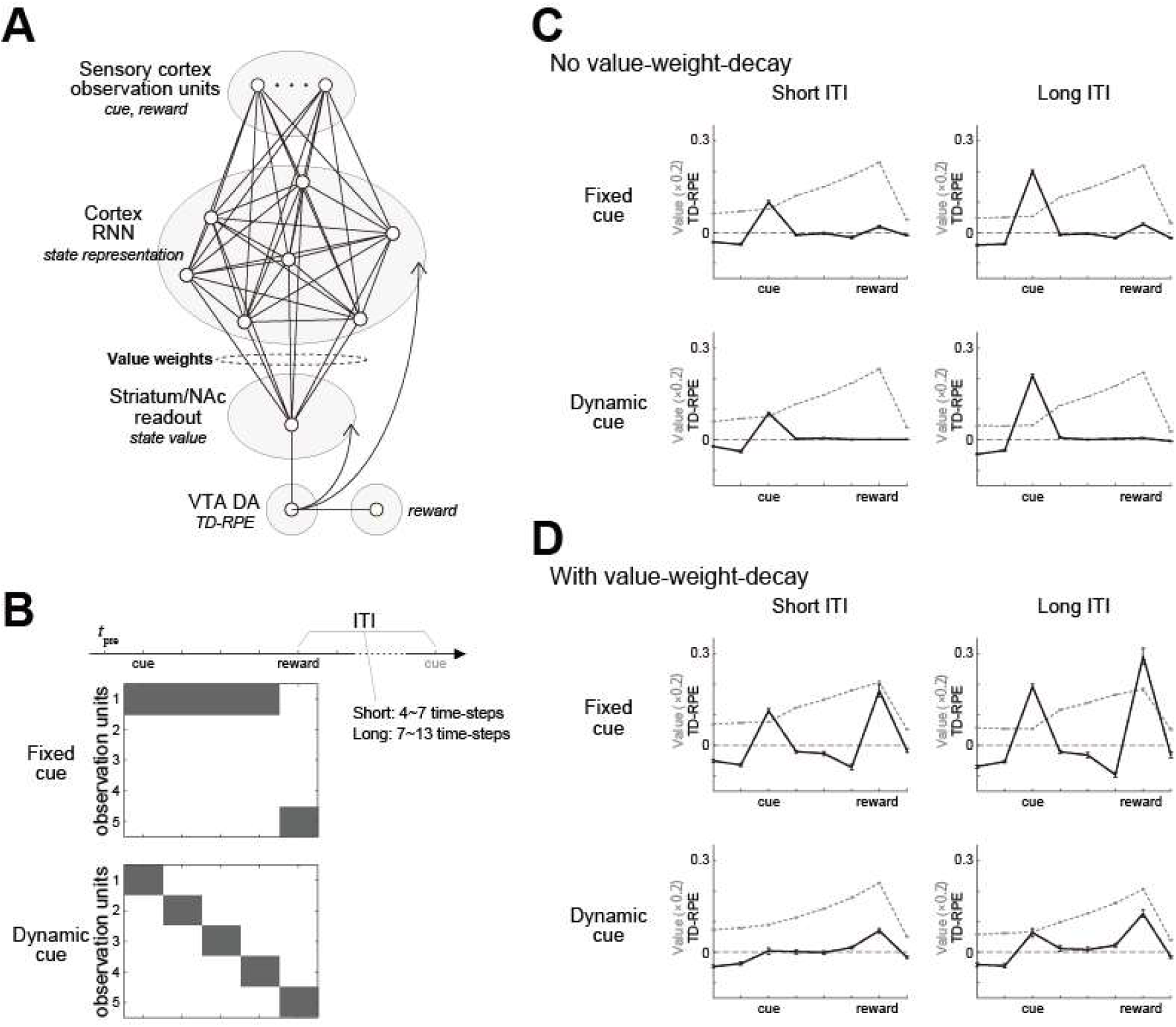
The online value-RNN model (oVRNN) with value-weight-decay generates different TD-RPE patterns depending on the cue type and the ITI length. **(A)** Schematic diagram of oVRNN, in which cortical RNN and its downstream striatum receive the same TD-RPE-encoding DA and learn state representation and value, respectively. **(B)** Simulation of cue-reward association task, varying the cue type and the ITI length. Fixed cue was simulated by that one of the observation units was continuously active during cue presentation, whereas dynamic cue was simulated by that different observation units sequentially became active. **(C**,**D)** Results when value-weight-decay was not assumed (C) or assumed (0.001 per time-step) (D). Black solid lines indicate TD-RPE in each condition. Gray dashed lines indicate state value, multiplied by a factor of five for display purposes. Error-bars indicate mean ± SEM across 100 simulations.

The observation units, whose activities at time-step *t* are denoted as ***o***(*t*) = (*o*_*k*_(*t*)), *k* = 1, …, 5), encode sensory information of cue (*o*_1_(*t*) ~ *o*_4_(*t*)) and reward (*o*_5_(*t*)), as detailed below. The RNN consists of recurrent units, whose activities (***x***(*t*) = (*x*_*j*_(*t*)), *j* = 1, ‥, 40) are determined by the activities of themselves and the observation units in the previous time-step:

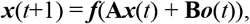

where **A** = (*A*_*ij*_) is the recurrent connection strength from *x*_*j*_ to *x*_*i*_ and **B** = (*B*_*ik*_) is the feed-forward connection strength from *o*_*k*_ to *x*_*i*_, and *f*(*z*) = 1/(1 + exp(−*z*)) is a sigmoidal function representing neuronal input-output relation. The activity of the readout unit *v*(*t*) is determined by:

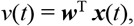

where ***w*** = (*w*_*j*_) are the value weights, presumably implemented by the cortico-striatal connection strengths. The TD-RPE unit encodes TD-RPE

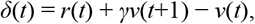

where *r*(*t*) is reward coming from the reward-encoding unit (*r*(*t*) = 1 at time-steps when reward was obtained, and *r*(*t*) = 0 at other time-steps), and *γ* is the time discount factor and was set to 0.8 in all the simulations shown in Figure 2.

The activities of RNN units (*x*_*j*_(*t*)) were initialized to pseudo uniform [0 1] random numbers. The elements of the value weights ***w*** were initialized to 0, and updated at every time-step as:

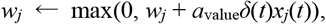

where max(*z*_1_, *z*_2_) returns the larger one of *z*_1_ and *z*_2_ (so that *w*_*j*_ was constrained to be non-negative) and *a*_value_ is the learning rate for the value weights and was set to 0.1/(40/12) = 0.03. Value-weight-decay was then applied at every time-step as:

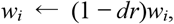

where *dr* was the decay rate per time-step and was set to 0 or 0.001.

The elements of the recurrent and feed-forward connection strengths **A** and **B** were initialized to pseudo standard normal random numbers, and updated at every time-step as:

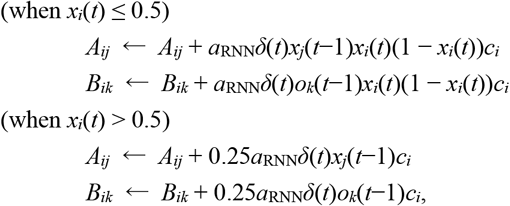

where *c*_*i*_ (*i* = 1, ‥, 40) were set to pseudo uniform [0 1] random numbers, and *a*_RNN_ is the learning rate for recurrent/feed-forward connections and was set to 0.1. This update rule was derived from the rule for the backpropagation method (backprop) with modifications for biological plausibility. Specifically, the symmetric feedback of backprop was replaced with the random feedback ***c*** = (*c*_*i*_) (c.f., (Lillicrap et al., 2016)), and the non-monotonic dependence on *x*_*i*_(*t*) (post-synaptic activity) was replaced with monotonic + saturation (as detailed in (Tsurumi et al., 2025)).

Using this oVRNN, we simulated cue-reward association tasks, with the type of cue (fixed or dynamic) and the length of inter-trial interval (ITI) (short or long) were varied (Fig. 2B). Presentation of a fixed cue was simulated by activating a single cue-corresponding observation unit for four consecutive time-steps, i.e., *o*_1_(*t*_pre_ + *k*) = 1 (*k* = 1, …, 4) where *t*_pre_ was the time-step immediately preceding cue presentation. In contrast, presentation of a dynamic cue was simulated by that the four cue-corresponding observation units became sequentially active, i.e., *o*_*k*_(*t*_pre_ + *k*) = 1 (*k* = 1, …, 4). For both types of cue, presentation/receival of reward was simulated by that reward-corresponding observation unit became active, i.e., *o*_5_(*t*_pre_ + 5) = 1. The observation units were nonactive (*o*_*k*_(*t*) = 0) in all the time-steps other than those mentioned above. ITI from the time-step of reward to the time-step of cue in the next trial was pseudo-randomly set to 4, 5, 6, or 7 time-steps in the short ITI conditions and 7, 9, 11, or 13 time-steps in the long ITI conditions. 100 simulations were conducted for each condition: there were in total 8 (= 2 × 2 × 2) conditions defined by the presence or absence of value-weight-decay, cue type (fixed or dynamic), and ITI length (short or long).

### Hierarchical RL model

We considered a hierarchical RL agent having coupled two learning systems, named the circuit M and circuit L, which modeled the CDh-rmSNc and CDt-clSNc circuits, respectively (Fig. 3A). We returned to the separate punctate representation of states. Each of these two circuits had the predicted value of each state, *V*_M_(*S*_*i*_) and *V*_L_(*S*_*i*_) (*i* = 1, …, *n*(= 5)) initialized to all 0. At each time-step/state, TD-RPE in each system was calculated as follows:

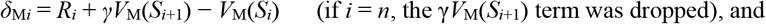

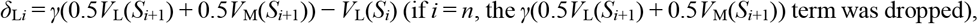

where *γ* was set to 1 in both (even if *γ* was instead set to 0.9, main points were largely preserved as shown in Extended Data Figure3-1). Notably, TD-RPE in the circuit L did not contain the reward term (*R*_*i*_) and instead contained the term for the predicted value of the upcoming state in the circuit M, *V*_M_(*S*_*i*+1_), which represented the conditioned reinforcement. Using these TD-RPEs, the predicted values in both circuits were updated as

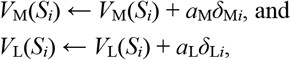

where *a*_M_ and *a*_L_ were the learning rates for the two circuits and were set as *a*_M_ = 0.25 or 0.125 (for Fig. 3F) and *a*_L_ = 0.025. At each time-step/state during the task trials, the predicted values of all the states decayed as

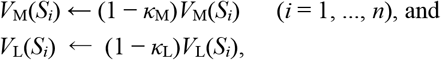

where *κ*_M_ and *κ*_L_ were the decay rates and were set as *κ*_M_ = 0.01 or 0.005 (for Fig. 3F) and *κ*_L_ = 0 (no decay) or 0.001 (for Fig. 3G,H). We also considered agent in which the − *V*_L_(*S*_*i*_) term in the calculation of *δ*_L*i*_ was replaced with − 0.5*V*_L_(*S*_*i*_) (Fig. 3C).

**Figure 3.**
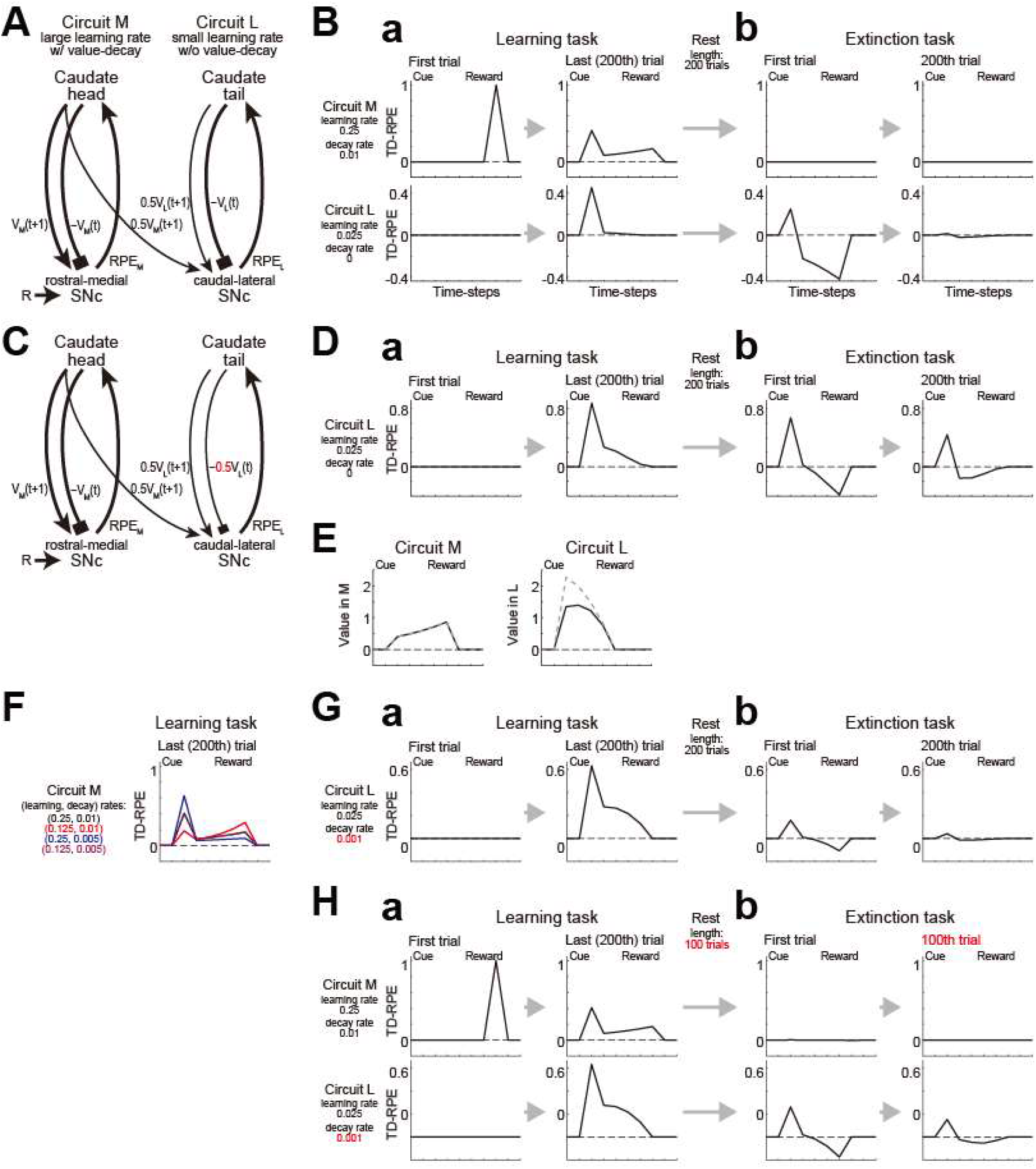
Hierarchical RL model of striatum-midbrain circuits with differences in learning rate and value-decay. **(A)** Hierarchical RL model. The circuit M, with a large learning rate (0.25) and value-decay (0.01), and the circuit L, with a small learning rate (0.025) and no value-decay, modeled the CDh-rmSNc and CDt-clSNc circuits, respectively. **(B)** TD-RPE generated in the model’s circuit M (top) and circuit L (bottom) during a simulated learning task (a) and a subsequent extinction task (b). Compare with the experimental results of DA neuronal activities in Figure 3A-E of (Kim et al., 2015). **(C)** The revised model with the strength of the negative “current state value” input in the circuit L reduced (halved: −0.5*V*_L_(*t*)), as indicated by the red color, so that it matched the strength of the “upcoming state value” input in the circuit L (0.5*V*_L_(*t*+1)) only, corresponding to the case where effective time discount factor (i.e., upcoming-value/current-value ratio) was 1 in the absence of the conditioned M→L input. **(D)** TD-RPE generated in the circuit L of the revised model. **(E)** State values in the circuit M (left) and circuit L (right) of the revised model at the last (200th) trial of the learning task (black solid lines). The gray dashed lines indicate the cases where the number of trials was increased to 400 (overlapped with the black solid line in the left panel). Compare with the experimental results of caudate neural activities in Figure 3B,C of (Kim and Hikosaka, 2013). **(F)** TD-RPE in the circuit M at the end of the learning task when either the learning rate or the decay rate was halved (red and blue lines, respectively) or both rates were halved (purple line) from the original case (black line). The purple and black lines are almost overlapped. **(G)** TD-RPE generated in the circuit L when value-decay of rate 0.001 was added to the circuit L so that the learning-rate/decay-rate ratio became common (=25) in the two circuits. **(H)** TD-RPE generated in the circuits M and L (learning rates: 0.25 and 0.025, decay rate: 0.01 and 0.001) when the lengths of the rest period and the extinction task were halved.

We simulated the behavior of this hierarchical RL agent in the cue-reward association learning task (200 trials, or 400 trials for Fig. 3E), and then continuously in the subsequent rest period and the extinction task. The rest period was assumed to have the same length as 200 trials or 100 trials (for Fig. 3H), and the predicted values of all the states decayed (if decay rate was not 0) 5 (number of time-steps/states) × 200 or 100 (number of trials) times according to the abovementioned equation. The extinction task (200 trials or 100 trials (for Fig. 3H)) was simulated in the same manner as the learning task except that the reward term (*R*_*i*_) was set to all 0.

### Distributional RL algorithms

We considered two distributional RL algorithms named the integRPE and segreRPE algorithms, which were extended from previously proposed models (Mikhael and Bogacz, 2016; Lowet et al., 2024). In both algorithms, there were two sets of predicted values of the states, *V*_1_(*S*_*i*_) and *V*_2_(*S*_*i*_) (*i* = 1, …, *n*), initialized to all 0. *V*_1_(*S*_*i*_) and *V*_2_(*S*_*i*_) were presumed to be represented by D1- and D2-receptor expressing direct and indirect-pathway striatal projection neurons (SPNs), respectively, and referred to as D1 and D2 values in the Results. In reality, D2 SPNs supposedly represent values in the sign-reversed manner (i.e., higher activity for lower predicted reward value), as suggested in experimental results (Lowet et al., 2024). Also, in order to allow changes of the values to both positive and negative directions under the constraint that the weights (strengths) of corticostriatal synapses should be non-negative, certain offsets of synaptic strengths / neural activity need to be presumed, in a similar manner to what we considered in our model for feature-based state representation in the above. However, here we did not model these biological details.

In the integRPE algorithm, at each time-step/state, a unified TD-RPE:

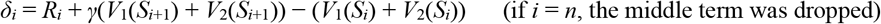

was calculated, where *γ* was set to 1. *V*_1_(*S*_*i*_) and *V*_2_(*S*_*i*_) were then updated as

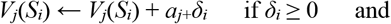

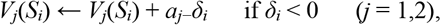

where the learning rates (*a*_1+_, *a*_1−_, *a*_2+_, *a*_2−_) were set to (0.15 0.05 0.05 0.15). At each time-step/state during the task trials, the two sets of predicted values of all the states decayed as

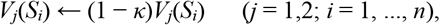

where *κ* was the decay rate and was set to 0 or 0.01. This algorithm is a TD version of the AU model (Mikhael and Bogacz, 2016).

In the segreRPE algorithm, at each time-step/state, segregated TD-RPEs:

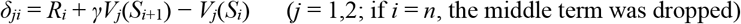

were calculated, where *γ* was set to 1. *V*_1_(*S*_*i*_) and *V*_2_(*S*_*i*_) were then updated as

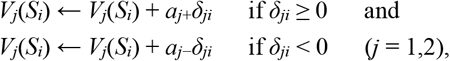

where the learning rates (*a*_1+_, *a*_1−_, *a*_2+_, *a*_2−_) were set to (0.15 0.05 0.05 0.15). Value-decay was not assumed in most simulations, except for those for Figure 4Bb-right, where value-decay (rate 0.01) was assumed in the same manner as in the integRPE algorithm. This segreRPE algorithm is a simplified TD version of the REDRL model (Lowet et al., 2024). The simplification that we made was to ignore the heterogeneities within D1 and D2 SPN populations. Specifically, in the REDRL simulation (Lowet et al., 2024), each of D1 and D2 populations consisted of five units with *a*_+_/(*a*_+_+*a*_−_) = 0.95, 0.85, …, 0.55 and 0.45, 0.35, …, 0.05, respectively, whereas in our segreRPE algorithm, single D1 and D2 populations had *a*_+_/(*a*_+_+*a*_−_) = 0.75 and 0.25, respectively. Also notably, the REDRL model first considered both asymmetric D1 and D2 plasticity for positive vs negative change in DA neuronal firing and asymmetric input-firing slope of individual DA neurons for positive vs negative RPE discovered experimentally (Dabney et al., 2020) and then combined these two asymmetries into integrated parameters (corresponding to *a*_+_ and *a*_−_ above). Therefore, in order to interpret our segreRPE algorithm as an simplification of the REDRL model, *a*_+_ and *a*_−_ should be similarly interpreted as integrated parameters, although in the Results we only mentioned the D1 vs D2 distinction in the introduction of this algorithm.

**Figure 4.**
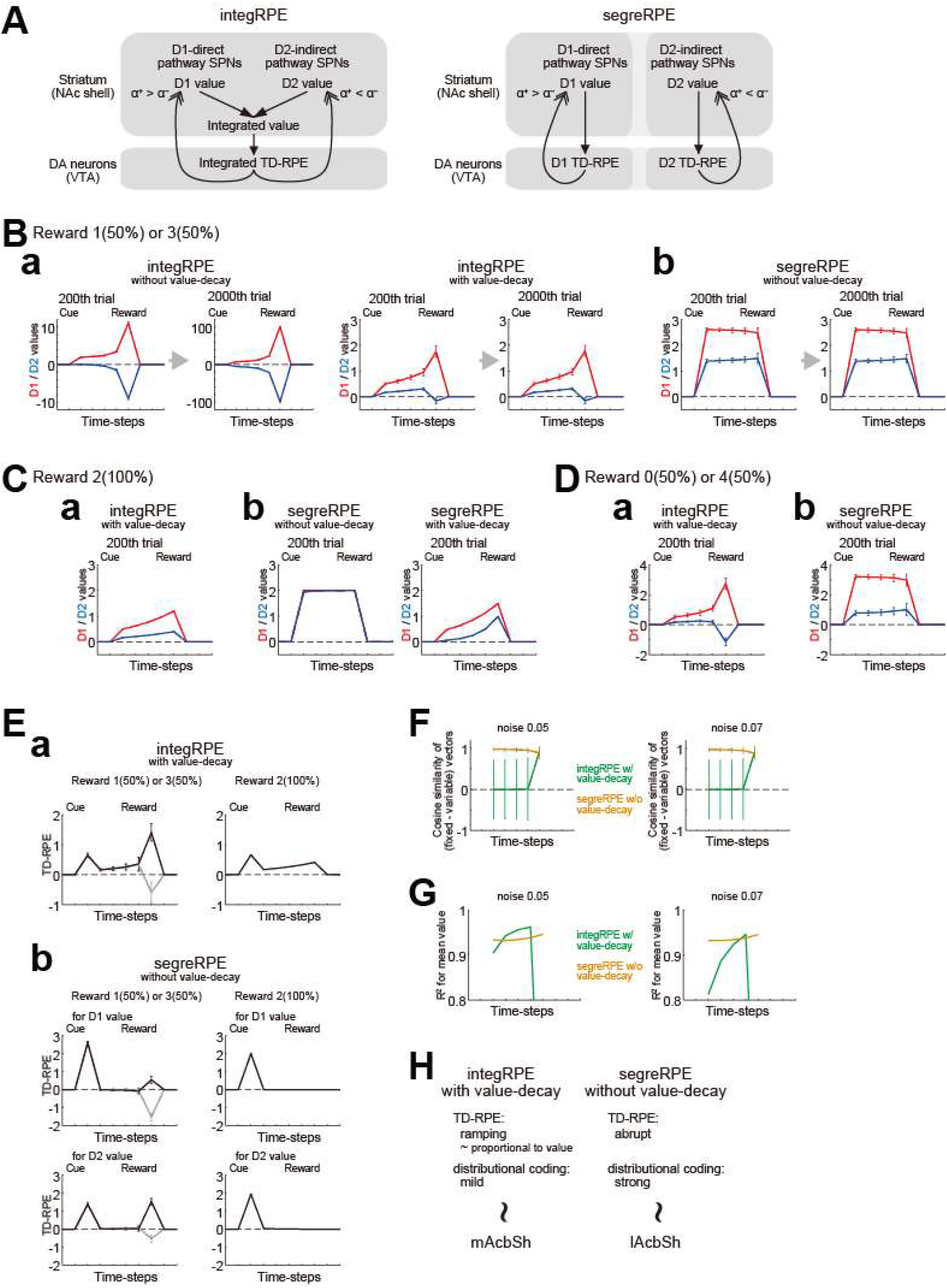
Behavior of different types of distributional RL algorithms with and without value-decay in probabilistic or deterministic cue-reward association learning tasks. **(A)** Different types of distributional RL algorithms. *Left*: integRPE, where D1 and D2 values are combined into the integrated value, from which the integrated TD-RPE is calculated and used for updates of both D1 and D2 values. *Right*: segreRPE, where D1 [/D2] TD-RPE is calculated only from D1 [/D2] values, and used only for updating D1 [/D2] value. **(B)** D1 values (red lines) and D2 values (blue lines) in a probabilistic task where reward was 1 or 3 with equal probabilities. The lines and error-bars indicate the mean±SD across 100 simulations (same applied to panels in (C,D)). **(a)** Results for the integRPE algorithm. Cases without value-decay (left two panels) and with value-decay (right two panels) at 200th and 2000th trials are shown. **(b)** Results for the segreRPE algorithm. Value-decay was not assumed, and cases at 200th and 2000th trials are shown. **(C)** D1 and D2 values in a deterministic task (reward: 2) in the integRPE algorithm with value-decay (a) and the segreRPE algorithm without value-decay (b-left: red and blue lines are almost overlapped) or with value-decay (b-right). **(D)** D1 and D2 values in a probabilistic task (reward: 0 or 4 with equal probabilities) in the integRPE algorithm with value-decay (a) and the segreRPE algorithm without value-decay (b). **(E)** TD-RPE in the probabilistic task (reward: 1 (gray lines) or 3 (black lines)) or in the deterministic task. The lines and error-bars indicate the mean±SD across simulations corresponding to each condition (reward size) within total 100 simulations. **(a)** Results for the integRPE algorithm with value-decay. **(b)** Results for the segreRPE algorithm without value-decay. Both TD-RPE for D1 values and TD-RPE for D2 values are shown. **(F)** Strength of reward distribution-coding quantified by an average cosine similarity of the difference in [D1 value, D2 value] vectors, at each time-step, between the condition where reward was always 2 (“fixed”) and the condition where reward was 1 or 3 with equal probabilities (“variable”). The green and orange lines indicate the integRPE algorithm with value-decay and the segreRPE algorithm without value-decay, respectively, and the lines and error-bars indicate the mean±SD across all possible pairs from 1000 simulations. **(G)** Strength of reward mean-coding quantified by *R*^2^ of correlation between the reward mean and the D1 value and D2 value, normalized across three conditions where reward was always 0, always 2, or 1 or 3 with equal probabilities. Note that the vertical axis starts at 0.8, and also that the result for the integRPE at the last (fifth) time-step was outside the drawing range in both panels. **(H)** Proposed correspondence between the types of distributional RL algorithms, the patterns of TD-RPE, the strengths of distributional coding, and the regions of AcbSh.

**Figure 5.**
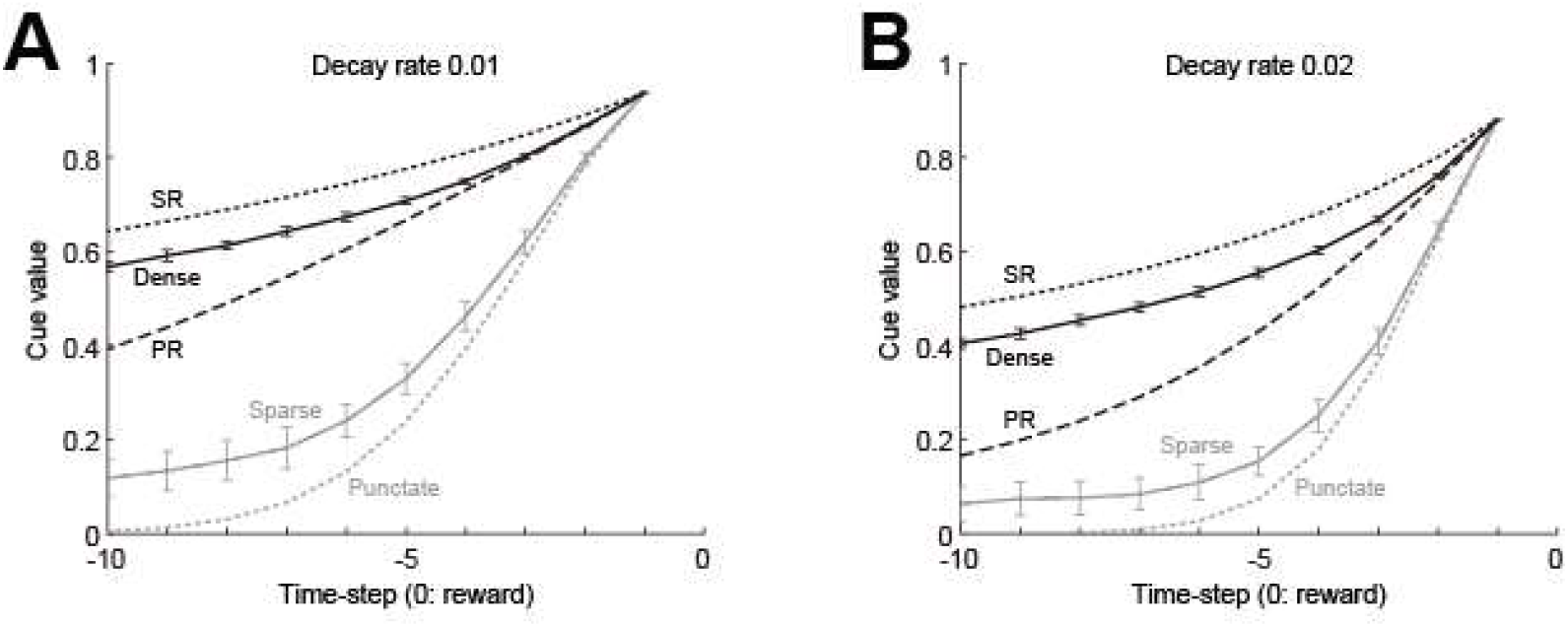
Dependence of effective temporal discounting on value-decay and state representation. The horizontal axis indicates the time-step of cue relative to reward (0), and the vertical axis indicates the cue value, i.e., TD-RPE upon cue. Decay rate was 0.01 (A) or 0.02 (B). The gray dotted, gray solid, black solid, black dotted, and black dashed lines indicate the cases with the punctate, sparse, dense, successor (SR), and predecessor (PR) representations, respectively. In the cases of sparse and dense representations, the lines and error-bars indicate the mean±SD across 100 simulations.

**Figure 6.**
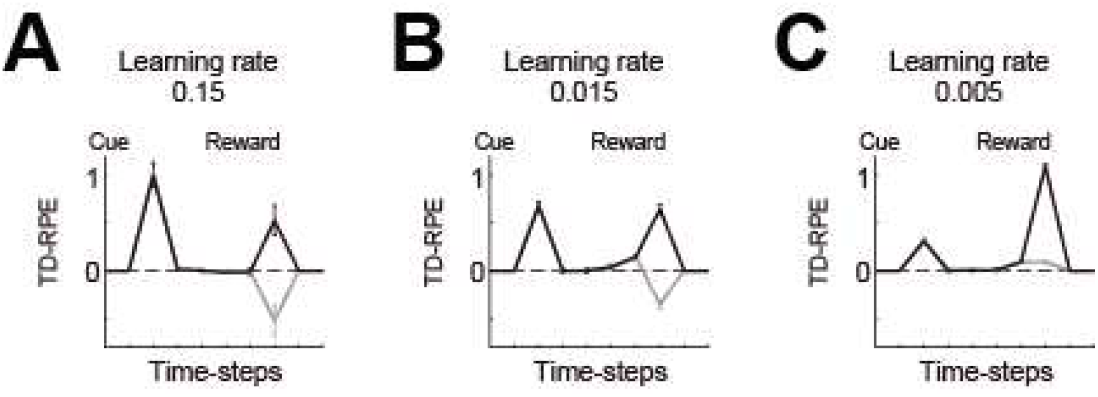
TD-RPE in a probabilistic cue-reward association task (reward: 0.5 (gray lines) or 1.5 (black lines) with equal probabilities) at 100th trial in the cases with different learning rates: 0.15 (A: learning is almost completed), 0.015 (B: learning is still incomplete), or 0.005 (C: learning is still incomplete). The lines and error-bars indicate the mean±SD across simulations corresponding to each condition (reward size) within total 100 simulations. Value-decay was not assumed, and the dense state representation (Fig. 1Ca) was assumed. Notably, if the states are instead represented in the punctate or sparse manner, in the period in which learning is incomplete, TD-RPE appears prominently at intermediate states between cue and reward and hardly or only limitedly at cue, deviating from the observed DA pattern in DLS (Tsutsui-Kimura et al., 2020).

We simulated the behavior of these algorithms in probabilistic or deterministic cue-reward association learning tasks (reward size: 1 or 3 with equal probabilities; 0 or 4 with equal probabilities; or 2 (100%)). Number of trials was set to 200 or 2000, and 100 simulations were conducted for each condition.

We quantified the strength of reward distribution- and mean-coding of the two algorithms in reference to the recent work (Lowet et al., 2024), through conducting a separate set of simulations assuming some noise. Specifically, for each of the integRPE algorithm with value-decay (decay rate 0.01) and the segreRPE algorithm without value-decay, we conducted 1000 simulations for each of three conditions (reward size: 0 (100%); 1 or 3 with equal probabilities; or 2 (100%)). We considered that the results of *i*-th simulations (*i* = 1, …, 1000) of the three conditions virtually correspond to the results of *i*-th pairwise recording from a D1 SPN and a D2 SPN in the three conditions (although in fact *i*-th simulations of the three conditions were just conducted independently and there was nothing in common). We repeated these entire simulations twice with assuming different levels of noise: pseudo-Gaussian random numbers of SD = 0.05 or 0.07 were added to D1 and D2 values at each time-step after these values for all time-steps were generated. As a measure of the strength of reward distribution-coding, we calculated an average cosine similarity of the difference in [D1 value, D2 value] vectors, at each time-step, between the condition where reward was always 2 (“fixed”) and the condition where reward was 1 or 3 with equal probabilities (“variable”). We calculated 1000 difference vectors using single simulations of the two conditions (i.e., *i*-th simulation of reward 2 − *i*-th simulation of reward 1 or 3 for *i* = 1, …, 1000) for each time-step. We then calculated the mean±SD of cosine similarity of each possible pair (in total 1000×999/2 = 499500 pairs) of these 1000 vectors (for each time-step). As a measure of the strength of reward mean-coding, we calculated *R*^2^ of correlation between the reward mean and normalized noise-added D1 value and normalized noise-added D2 value (normalized, separately for D1 value and D2 value, across the three conditions for each of 1000 simulations).

Even if the time discount factor *γ*, which was set to 1 in both integRPE and segreRPE algorithms, was instead set to 0.9 in both algorithms, main points were largely preserved as shown in Extended Data Figure4-1.

### Software

Simulations were conducted by using MATLAB, and pseudo-random numbers were implemented by using the functions rand, randn, and randperm.

### Code Accessibility

The codes used in the present work will be deposited in GitHub upon acceptance and publication of this manuscript.

## Results

### Value-decay in RL under different state representations

We examined the pattern of TD-RPE generated in RL models with value-decay under different state representations in a simulated cue-reward Pavlovian association learning task, in which a single cue was followed by a single reward of a fixed amount after a fixed delay (see the Methods for the details of the models and the simulations). First we examined a case where discretized individual timings from cue to reward were represented separately as punctate states (Fig. 1Aa), as done previously (Morita and Kato, 2014). The value of state *S*_*i*_, *V*(*S*_*i*_), was updated by TD-RPE, *δ*_*i*_ = *R*_*i*_ + *γV*(*S*_*i*+1_) − *V*(*S*_*i*_), where *R*_*i*_ was obtained reward, as *V*(*S*_*i*_) ← *V*(*S*_*i*_) + *aδ*_*i*_, where *a* was the learning rate, and value-decay was implemented at each time-step/state as *V*(*S*_*i*_) ← (1 − *κ*)*V*(*S*_*i*_), where *κ* was the decay rate (see the Methods for details). In this case, if there was value-decay, TD-RPE showed a sustained ramp towards reward after learning, in addition to an abrupt response to cue (Fig. 1Ac). As the rate of value-decay increased, ramping became more prominent, while the abrupt response to cue degraded (Fig. 1Ad). Notably, the effect of value-decay on the pattern of TD-RPE is different from the effect of temporal discounting. Specifically, value-decay causes diminished abrupt TD-RPE upon cue and ramping TD-RPE toward reward whereas temporal discounting causes only diminished abrupt TD-RPE upon cue, while both value-decay and temporal discounting cause ramping of state values (Fig. 1F).

Next we examined cases where the states were represented by features. Specifically, we assumed that each state *S*_*i*_ was represented by a high-dimensional feature vector, ***x***_*i*_ = (*x*_*i*,1_, …, *x*_*i*,100_), and state values were calculated as weighted sums of features with subtraction of constants: *V*(*S*_*i*_) = ***w***·***x***_*i*_ − ***w***_**0**_·***x***_*i*_, where ***w*** = (*w*_1_, …, *w*_*m*_) was the weight vector and ***w***_**0**_ = (*w*_0_, …, *w*_0_) was a constant vector to which ***w*** was initialized. The weights were updated according to TD-RPE multiplied with features (Sutton and Barto, 2018). Biologically, features were assumed to be encoded by cortical neurons and the update was assumed to be implemented by DA-dependent plasticity of corticostriatal synapses (Montague et al., 1996; Sutton and Barto, 2018). We further assumed non-negativity of each weight for biological plausibility, although negative weights were effectively allowed by the assumed subtraction of constants in the value calculation (presumably implemented by striatal inhibitory interneurons). We also assumed a larger learning rate for positive than negative TD-RPE, in reference to findings and suggestions on the property of D1 receptor-dependent plasticity (Collins and Frank, 2014; Mikhael and Bogacz, 2016; Lee et al., 2021; Pinto and Uchida, 2023; Lowet et al., 2024), resulting in the update rule: ***w*** ← max(0, ***w*** + *a*_f_*δ*_*i*_***x***_*i*_/||***x***_*i*_||^2^), where the learning rate *a*_f_ was *a*_+_ if *δ*_*i*_ ≥ 0 and *a*_−_ if *δ*_*i*_ < 0 and *a*_+_ > *a*_−_ was assumed. Value-decay (value-weight-decay) was implemented at each time-step/state as *w*_*i*_ ← *w*_0_ + (1 − *κ*)(*w*_*i*_ − *w*_0_).

Since it has been suggested that cortical representations may be sparse (Barth and Poulet, 2012), we examined a case where, for each feature vector, only a small proportion (10%) of elements had positive pseudo-random values (Fig. 1Ba). As the rate of decay of value-weights (to the initial weights) increased, TD-RPE exhibited a ramp towards reward (Fig. 1Bc,d), similarly to the case with punctate representation. We also examined a case with dense representation, where all the elements of each feature vector had positive pseudo-random values (Fig. 1Ca). In this case, value-weight-decay still caused ramping of TD-RPE, but its degree was smaller than the case with sparse representation (Fig. 1Cc,d).

We further examined a case where the states were represented by future occupancies of other (and self) states, namely, by the successor representation (SR) (Dayan, 1993) (Fig. 1Da), which enables partially model-based-like behavior (Russek et al., 2017; Stachenfeld et al., 2017) and has been suggested to be actually used by humans (Momennejad et al., 2017) and implemented in certain parts of cortico-striatal-DA system (Garvert et al., 2017; Russek et al., 2021). In this case, value-weight-decay did not cause a ramp of TD-RPE while still causing sustained response to reward (Fig. 1Dc,d). We considered that the absence of TD-RPE ramping in the case with SR could provide an explanation of the observed gradual fading of DA ramping over daily training of reward navigation (Guru et al., 2020). Indeed, we found that when representation of states towards reward was initially punctate but gradually became closer to SR through learning of SR features, TD-RPE ramping appeared in initial/early trails but then gradually faded away (Fig. 1Gc, compare to Fig. 4a of (Guru et al., 2020)). Notably, TD-RPE prominently ramped at a later (reward-proximal) phase in initial trials (Fig. 1Gb) but more constantly ramped in subsequent trials (Fig. 1Ga), resembling the observed patterns of DA ramping (Fig. 1j and 1e of (Guru et al., 2020)).

As so far shown, value(−weight)-decay generally caused sustained TD-RPE at reward after learning completion, and also ramping TD-RPE towards reward except for the case with SR, but their degrees heavily depended on state representation. Given that dense representation, in which feature vectors were much overlapped, caused small ramping and SR, in which feature vectors are nested and so in a sense completely overlapped, caused no ramping, it could be guessed that the degrees of ramping depend, negatively, on the degree of overlaps between feature vectors. To test this, we examined a case with ‘predecessor’ representation (PR) (c.f., (Bailey and Mattar, 2022)), where the states were represented by past, rather than future, occupancies of other (and self) states. In this case, value-weight-decay caused rather small sustained TD-RPE at reward and little ramping-up but instead ramping-down (Fig. 1E). Because the PR feature vectors had the same degree of overlaps as the SR feature vectors had, this result suggests that not only the degree of overlaps between feature vectors but also their temporal order affects the pattern of TD-RPE.

So far we assumed that cortical state representation is fixed, and not affected by TD-RPE-encoding DA signals. However, subpopulation of VTA DA neurons project also to the cortex, especially, its frontal regions (Williams and Goldman-Rakic, 1998), and it has been suggested that cortical DA, as well as striatal DA, encodes (TD-)RPE (O’Doherty et al., 2003; Takahashi et al., 2011; Starkweather et al., 2018) and modulates plasticity (Otani et al., 2003). Computational role of TD-RPE-encoding cortical DA remained unclear, but recent modeling work (Tsurumi et al., 2025), extending previous studies (Hennig et al., 2023; Qian et al., 2025), suggested its role in learning of state representation appropriate for task. Specifically, that model, named the online value-RNN (oVRNN), suggested that if cortical recurrent neural network (RNN) and its downstream striatum receive the same TD-RPE-encoding DA, state representation and value can be simultaneously learned in the cortex and striatum, respectively (Fig. 2A). Moreover, value-weight-decay in oVRNN was shown to promote ‘alignment’ of the value weights (corticostriatal projections) and the feedback weights (mesocortical projections), which is potentially useful for learning (Lillicrap et al., 2016), while the effect of value-weight-decay on the pattern of TD-RPE was not examined.

In oVRNN, learning was assumed to be continuous across trials, including inter-trial-intervals (ITIs). Recent work (Floeder et al., 2024) found that whether DA in NAc core exhibits a ramp or not depends on the length of ITI and the nature of cue. Specifically, DA ramped when ITI was short and the cue was dynamic (i.e., changing tone), but not when ITI was long or the cue was fixed (fixed tone). We examined if oVRNN, with value-weight-decay, could explain this finding through simulations with varying cue type and ITI length (Fig. 2B). When there was no value-weight-decay, TD-RPE showed a prominent peak at cue and no ramp regardless of cue type or ITI length (Fig. 2C). When value-weight-decay was introduced, if the cue was dynamic and ITI was short, TD-RPE did not show a prominent peak at cue, and instead showed an overall increasing pattern, although pre-reward ramping was only slight (Fig. 2D, bottom-left). In contrast, if the cue was fixed or ITI was long, TD-RPE showed a prominent peak at cue, although slight pre-reward ramping appeared in the dynamic & long condition. These results roughly resemble the experimental finding (Floeder et al., 2024). The cue-type-dependance of the TD-RPE pattern is considered to be because representations of different time-steps would become less overlapped when the cue is dynamic rather than fixed.

### Hierarchical RL model consisting of circuits with and without value-decay

As mentioned in the Introduction, it was shown that CDh neurons rapidly learned the cue value within a session but forgot it in subsequent days while CDt neurons gradually developed stable value-memory (Kim and Hikosaka, 2013). Also, DA neurons in rostral-medial substantia nigra pars compacta (rmSNc), which projects to CDh, forgot predicted-value-memory whereas DA neurons in caudal-lateral SNc (clSNc), which projects to CDt, retained it (Kim et al., 2015). Specifically, rmSNc DA neurons developed TD-RPE response during the cue-reward association task but had lost it at the subsequent passive viewing task, whereas clSNc DA neurons retained TD-RPE response in the latter task. These results appear to be potentially explained if CDh-rmSNc circuit implements RL with a large learning rate and value-decay whereas CDt-clSNc circuit implements RL with a small learning rate and no value-decay. But how do these two circuit interact? It was found (Kim et al., 2015) that the memory-retain type clSNc DA neurons did not respond to unpredicted reward. The authors suggested that these DA neurons may instead receive ‘conditioned reinforcement’ information from the forget/update type rmSNc DA neurons, which responded to unpredicted reward, potentially through the superior colliculus (Kim et al., 2015). Such conditioned reinforcement information might also be sent through the suggested ‘spiral’ structure of the striatum-midbrain circuits (Haber et al., 2000; Yin et al., 2008).

Based on these findings and considerations, we constructed a hierarchical RL model (c.f., (Haruno and Kawato, 2006; Ito and Doya, 2010; Botvinick, 2012; Frank and Badre, 2012; Keramati and Gutkin, 2013; Balleine et al., 2015)) consisting of coupled two circuits, the circuit M, modeling CDh-rmSNc, and the circuit L, modeling CDt-clSNc (Fig. 3A). The circuit M had a (relatively) large learning rate and value-decay whereas the circuit L had a small learning rate and no value-decay. Also, the circuit M receives external reward information, whereas circuit L does not receive direct reward input but receives the information of upcoming state value from the circuit M. We simulated the cue-reward association learning task by this model, assuming the punctate state representation. As learning progressed, TD-RPEs in both circuits M and L responded to cue while TD-RPE in the circuit M additionally showed sustained response to reward (Fig. 3Ba). These patterns look similar to the reported DA patterns in clSNc and rmSNc (Figure 3A-E of (Kim et al., 2015)) including a small apparent response of rmSNc DA to reward although it could just reflect contributions of initial trials.

In the experiment (Kim et al., 2015), after the learning task, monkeys engaged in free viewing and passive viewing sessions, in which multiple stimuli were simultaneously or sequentially presented without contingent reward, and memory retention was examined. In our model, after the learning task (200 trials), we simulated a rest period with the same duration (corresponding to 200 trials), and then an extinction task (200 trials), in which the learned cue was presented without reward. At the beginning of the extinction task, TD-RPE in the circuit M disappeared whereas TD-RPE upon cue in the circuit L was retained (Fig. 3Bb-left), in line with the disappearance and retention of rmSNc and clSNc neuronal response, respectively (Kim et al., 2015). However, the positive cue response of TD-RPE in the circuit L was followed by a large negative response (Fig. 3Bb-left-bottom) and these responses almost disappeared after extinction learning (Fig. 3Bb-right-bottom), deviating from the experimental result.

We explored how this discrepancy could be mitigated, and found that reducing the coefficient of the negative “current state value” input (− *V*_L_(*S*_*i*_)) in the circuit L (Fig. 3C: where the coefficient was halved (from 1 to 0.5)) was effective, as shown in Figure 3D, although some negative response still remained. In the original model (Fig. 3A), this coefficient was set to match the sum of coefficients of the “upcoming state value” inputs from the circuit M (0.5*V*_M_(*S*_*i*+1_)) and the circuit L itself (0.5*V*_L_(*S*_*i*+1_)) (i.e., 0.5 + 0.5 = 1), corresponding to the case where effective time discount factor (i.e., upcoming-value/current-value ratio) was 1 in the presence of the conditioned M→L input. The revised model (Fig. 3C) instead corresponded to the case where effective time discount factor was 1 in the absence of the M→L input. The better reproduction of the experimentally observed long-term retention of clSNc DA response by the latter model indicates that the actual striatum-DA circuits are tuned in that way: in the extinction task, in the circuit L, the magnitude of the negative “current value” input is not much larger than the magnitude of the positive “upcoming value” input so that TD-RPE does not become quite negative, ensuring long-term retention of response.

If that is the case, in the presence of the M→L input, effective time discount factor exceeded 1 in the circuit L, and so delay should cause amplification, rather than discounting, of reward value. Indeed, the value in the circuit L at the end of the learning task showed such a pattern, i.e., backwardly ramped towards the early phase of object presentation (Fig. 3E-right, black solid line). By contrast, the value in the circuit M showed a ramp towards the late phase, reflecting temporal discounting caused by value-decay (Fig. 3E-left, black solid line). When the number of trials of the learning task was increased, the cue response in the circuit L was further increased while the activity of the circuit M remained unchanged (Fig. 3E, gray dashed lines). Crucially, these temporally increasing and decreasing values in the circuits M and L appear to match the observed patterns of CDh and CDt neuronal activities, respectively (Figure 3B,C of (Kim and Hikosaka, 2013)), providing a possible mechanistic explanation. Functionally, such a value amplification in the circuit L is considered to contribute to long-term value retention after reward extinction, forming a stable habit while potentially causing a risk of addiction.

So for we regarded the rates of value learning and decay as independent parameters. However, learning and decay might be interlinked at the level of physiological mechanisms, and if so, their rates may co-vary. Figure 3F illustrates how varying the rates of learning and decay affects TD-RPE in the circuit M at the end of the learning task. When either the learning rate or the decay rate was halved, the TD-RPE pattern changed noticeably (red and blue lines in Fig. 3F). In contrast, when both rates were halved, the pattern remained almost unchanged (purple line), implying a potential importance of the learning/decay ratio. Given these considerations, we asked if the observed DA patterns could still be reproduced when assuming that the circuits M and L have the same ratio of learning/decay. Specifically, we examined a modified case where the circuit L, which was originally decay-free, was re-assumed to have value-decay so that its learning/decay ratio became equal to that of the circuit M. As a result, TD-RPE in the circuit L became too small at the end of the extinction task (Fig. 3G). Nonetheless, this issue was mitigated when the lengths of the rest period and the extinction task were halved (Fig. 3H), Therefore, our decay-based account of the neural activity patterns could be in line with, though not especially consistent with, the possible covariation of learning and decay.

### Distributional RL algorithms with and without value-decay

Recent studies suggested that the cortico-basal ganglia-DA system implements distributional RL (Mikhael and Bogacz, 2016; Dabney et al., 2020; Lowet et al., 2020; Pinto and Uchida, 2023; Lowet et al., 2024; Muller et al., 2024). Given that D1-direct-pathway and D2-indirect-pathway striatal projection neurons (SPNs) predominantly learn from positive and negative TD-RPE, respectively (c.f. (Collins and Frank, 2014; Mikhael and Bogacz, 2016; Iino et al., 2020; Lee et al., 2021)), they can develop positively biased D1 value and negatively biased D2 value (Mikhael and Bogacz, 2016) and thus collectively have information of reward distribution. There are two (extreme) possibilities on how TD-RPE is calculated from D1 and D2 values. The first one is that D1 and D2 values are combined (summed) into an integrated value, from which an integrated TD-RPE is calculated and used for updates of both D1 and D2 values. We refer to this as the integRPE algorithm (Fig. 4A left), which is a TD version (extension) of the Actor learning Uncertainty (AU) model (Mikhael and Bogacz, 2016). The second possibility is that pathway-specific TD-RPEs are calculated based on either D1 value or D2 value only, and each pathway is updated using such a segregated TD-RPE. We refer to this as the segreRPE algorithm (Fig. 4A right), which is a TD version (extension), with simplifications, of the reflected EDRL (REDRL) model (Lowet et al., 2024) (an extension of the Expectile Distributional RL (EDRL) (Rowland et al., 2019)).

We simulated a probabilistic cue-reward association learning task, in which reward of either size 1 or 3 was obtained with equal probabilities, using each of these two algorithms. When the integRPE algorithm was used, D1 values became positive whereas D2 values became mostly negative (Fig. 4Ba-leftmost), and the magnitudes of both of these values grew unboundedly (compare the values at 2000th trial with those at 200th trial in Fig. 4Ba-left), while the magnitudes of the integrated value and TD-RPE did not. Such unbounded value growth, which seems biologically implausible, was prevented if value-decay was incorporated (Fig. 4Ba-right). In this sense, value-decay is necessary for the integRPE algorithm, and indeed value-decay was assumed in the original AU model (Mikhael and Bogacz, 2016). In contrast, when the segreRPE algorithm was used, both D1 and D2 values became positive, and they did not grow unboundedly even without value-decay (Fig. 4Bb). Thus, value-decay is not necessary for the segreRPE algorithm, and it was not assumed in the original REDRL model (Lowet et al., 2024).

We also simulated a deterministic cue-reward association task, in which reward of size 2 (equal to the mean of the reward sizes in the abovementioned probabilistic task) was always obtained, again using each of the two algorithms. When the integRPE algorithm (with value-decay) was used, D1 and D2 values became both positive but separated; D1 value became larger than D2 value (Fig. 4Ca). In contrast, when the segreRPE algorithm (without value-decay) was used, D1 and D2 values converged to almost the same positive value (Fig. 4Cb-left). This way, the segreRPE algorithm better discriminated the probabilistic reward and the deterministic reward than the integRPE algorithm (although the integRPE algorithm could still encode the reward split (c.f. (Mikhael and Bogacz, 2016)) as apparent from the result for a task with a larger reward split (0 or 4) (Fig. 4Db)). Notably, if value-decay was assumed in the segreRPE algorithm, D1 and D2 values no longer converged to an almost same value but to different values (Fig. 4Cb-right). Thus, value-decay is not only unnecessary but harmful for the segreRPE algorithm to achieve the ideal differential coding of the different reward distributions.

The integRPE algorithm with value-decay generated ramping TD-RPE (Fig. 4Ea) whereas the segreRPE algorithm without value-decay generated only abrupt TD-RPE (Fig. 4Eb). Also, in the integRPE algorithm with value-decay, D1 value ramped up and D2 values also (less prominently) ramped up except for the last part, whereas D1 and D2 values are largely flat in the segreRPE algorithm without value-decay (Fig. 4B-D). Regarding the coding property, the segreRPE algorithm without value-decay better discriminated the different reward distributions, and so could realize better distribution-coding than the integRPE algorithm with value-decay. Conversely, the integRPE algorithm with value-decay encoded different distributions with the same mean more similarly, and so could realize better mean-coding than the segreRPE algorithm without value-decay. We quantified the strength of distribution- and mean-coding of the two algorithms (c.f., (Lowet et al., 2024)), through simulations assuming some noise (see the Methods for details). The results confirmed and refined the abovementioned conjectures. As for the strength of distribution-coding (Fig. 4F), the score of the segreRPE algorithm without value-decay was high for all the time-steps from cue to reward whereas the score of the integRPE algorithm with value-decay was low except for the last time-step, regardless of the noise levels. Regarding the strength of reward mean-coding (Fig. 4G), while the score of the segreRPE algorithm without value-decay was nearly constant across time-steps, the score of the integRPE algorithm with value-decay showed a gradual increase, surpassing the segreRPE when the noise was small (Fig. 4G-left), and a drop at the last time-step. The gradual increase in the integRPE’s mean coding reflects the increase (ramp) of the D1/D2 values and a resulting improvement in the signal-to-noise ratio.

Given these differential properties, here we propose that the integRPE and segreRPE algorithms can coherently explain recent findings on DA patterns and coding properties in the NAc shell. As for the DA pattern, recent work (de Jong et al., 2024) found that DA in lateral NAc shell (lAcbSh) showed an abrupt cue response whereas DA in medial NAc shell (mAcbSh) showed a sustained/ramping pattern. These DA patterns appear to be consistent with a previous finding (Kim et al., 2020) that medial VTA DA neurons exhibited more positive ramps than lateral VTA DA neurons. Regarding the coding property, another recent work (Lowet et al., 2024) showed that distributional coding was strongest in lAcbSh and weaker in mAcbSh (although *n* = 1 for mAcbSh). As for mean coding, the strength in lAcbSh was relatively constant between cue and pre-reward timings whereas the strength in mAcbSh ramped and surpassed that of lAcbSh (Lowet et al., 2024). We argue that these results for DA patterns and AcbSh coding properties can be coherently explained if lAcbSh and mAcbSh implement the segreRPE algorithm without value-decay and the integRPE algorithm with value-decay, respectively (Fig. 4H). Crucially, this explanation can reconcile the controversy on whether ramping DA in mAcbSh/medial VTA represents value (de Jong et al., 2024) or TD-RPE (Kim et al., 2020). Specifically, because ramping TD-RPEs in the case with value-decay can be proportional to state values, except for cue response, according to the formulae that we previously derived (Morita and Kato, 2014), the ramping DA in mAcbSh could in fact represent both TD-RPE and value.

## Discussion

We have shown how value-decay links heterogeneous DA patterns and various RL computations: (i) whether value-decay causes ramping TD-RPE depends on state representation, potentially explaining condition-dependent appearance/fading of DA ramping, (ii) the hierarchical RL model consisting of circuits with and without value-decay explains regional differences in DA/striatal activity patterns and memory flexibility/stability, and (iii) the distributional RL algorithms with and without value-decay coherently explain regional differences in DA patterns and strengths of distributional coding. Below we discuss each of these.

Our model uniquely predicts that the degree of over-trial/session retention of DA/striatal response is associated with within-trial temporal pattern, i.e., forgetful DA shows sustained/ramping patterns and stably retained DA shows abrupt patterns, and we further predict a medial-lateral gradient. A previous result in (Iino et al., 2020) could be consistent with this prediction, though not tested. Specifically, DA terminal response to reward-associated cue remained on the next day of training in lateral NAc core whereas it disappeared in medial NAc core, and on the last day of training, lateral DA showed abrupt response to cue whereas medial DA showed a small ramp toward reward (their Extended Data Fig. 3f,g). Medial DA did not ramp on preceding days, but it could be because learning of state representation (as in our simulation in Fig. 2) took time while sensory-driven representation was used in the lateral side. These points were not focused in that study, and may be difficult to examine because cue-reward delay was short. Therefore, these are desired to be tested in future experiment with longer cue-reward delay and systematic examination of response retention and patterns.

### Value-decay and state representation

Our results suggested that gradual fading of DA ramping as training progressed in reward navigation (Guru et al., 2020) could result from gradual formation of SR. Persistence of DA ramping in another wheel-running task without spatial cues in the same study (Guru et al., 2020) could also be explained by no formation of SR due to the lack of spatial cues. The authors suggested that the ramping DA was not RPE because it quickly appeared in the next trial after the first reward was obtained. But quick RPE-like DA response was reported previously (Kim et al., 2015), and the observation that DA ramping appeared in a late timing in initial trials but appeared from the beginning in later trials (Fig. 1j and 1e of (Guru et al., 2020)) matches the property of TD-RPE in our simulations (Fig. 1Ga,b). A limitation of our present model/simulation is that passive state transition and active transition by voluntary action (Hamid et al., 2021), as well as state-value/critic learning and action-value/actor learning (Joel et al., 2002; O’Doherty et al., 2004; Takahashi et al., 2008; Sutton and Barto, 2018; Fraser et al., 2023), are not distinguished. Incorporation of them would be an important future direction.

Value-decay caused a decrease of TD-RPE upon cue. This can be regarded as effective temporal discounting of cue value (Kato and Morita, 2016), which is distinct from genuine temporal discounting (time discount factor <1) as we have shown (Fig. 1F). Our simulations suggest that the degree of effective temporal discounting by value-decay depends on state representation (Fig. 5). Given these, caution would be needed when estimating the time discount factor from DA patterns (Masset et al., 2023; Mohebi et al., 2024).

### Hierarchical RL with different learning and value-decay rates

In the experiments, caudate and DA neurons showed response not only to stimuli associated with reward but also, to lesser extents in most cases, to those associated with no reward. Such responses are not explained by our present model. They could potentially relate to generalization or context (Mirenowicz and Schultz, 1996; Kobayashi and Schultz, 2014; Matsumoto et al., 2016), and a possible future direction is to consider models incorporating feature-based representation (Lee et al., 2024) that enables generalization. Response to no-reward stimuli in CDt-clSNc could also reflect saliency. CDh and CDt have homologies to rodent DMS (Balleine and O’Doherty, 2010) and tail-of-striatum (TS) (Jiang and Kim, 2018; Lee et al., 2023; Green et al., 2024), respectively, although CDt has stronger vision-selectivity than TS (Jiang and Kim, 2018; Lee et al., 2023). Stable-value-encoding in CDt (Kim and Hikosaka, 2013; Kim et al., 2015) and potential-threat-encoding in TS (Menegas et al., 2018; Akiti et al., 2022; Tsutsui-Kimura et al., 2025) could then be commonly understood as saliency encoding (Green et al., 2024).

Along with CDt/TS, rodent dorsolateral striatum (DLS) was shown to be crucial for stimulus-response behavior (Yin et al., 2004). Since DA in DLS responds to unpredicted reward, different from TS, DLS could be considered as an intermediate between DMS and TS. Then, our model predicts smaller learning rate and value-decay in DLS than in DMS. In cue-reward association task with multiple reward sizes (Tsutsui-Kimura et al., 2020), DA in DMS showed positive and negative response to large and small reward, respectively, but DA in DLS showed positive response even to small reward. This could be because the learning rate was small and thereby learning was still incomplete (Fig. 6). Possibly, the rates of learning and value-decay are gradually distributed from DMS to DLS (and to TS). It may explain a larger variety of DA patterns. In particular, resulting continuous spectrum of abrupt and ramping patterns could manifest as spatial waves, potentially underlying observed medial→lateral and lateral→medial DA waves (Hamid et al., 2021).

A recent study (Mohebi et al., 2024) suggested that DA dynamics is fast in DLS, slower in DMS, and even slower in the ventral striatum (VS). At first glance, this looks contradictory to our proposal of smaller learning rate and value-decay in DLS than in DMS. However, these two might be the opposite sides of the same coin. DLS may steadily learn the value of individual short action-elements (movements), and DA/RPE there is abrupt since there is no value-decay (value decreases due to negative RPE). In contrast, DMS (and VS) may quickly learn and forget the value of macro-level actions, which reflect the context, such as the current reward richness/poorness, coming from the environmental model and can thus fluctuate in a slow time scale.

### Distributional RL algorithms with and without value-decay

We have presented two algorithms of distributional RL, integRPE and segreRPE, summarizing and extending the previous models (Mikhael and Bogacz, 2016; Pinto and Uchida, 2023; Lowet et al., 2024), and shown that their implementations in different striatal regions, mAcbSh and lAcbSh, respectively, can coherently explain the regional differences in DA patterns (de Jong et al., 2024) and the strengths of distributional coding (Lowet et al., 2024). Note, however, that the result for coding strength is currently based on limited sample (*n* = 1 for mAcbSh) (Lowet et al., 2024), and so it could rather be regarded as a prediction that awaits further experiment.

Given that the segreRPE algorithm achieves better distributional coding, what can the integRPE algorithm be beneficial for? One possibility is an anatomical merit, or rather a constraint. The segreRPE algorithm requires that segregated RPEs for D1 and D2 values are calculated and utilized without intermixing, but it may be difficult because of the widespread dense axonal projections of DA neurons (Matsuda et al., 2009) and intermingled placements of D1 and D2 SPNs in most striatal regions albeit with exceptions (Gangarossa et al., 2013; Murata et al., 2015; Petryszyn et al., 2017; Ogata et al., 2022). The integRPE algorithm can be more simply implemented through broadcasting DA signals.

The integRPE algorithm might also have functional merits. Because individual DA neurons encode the integrated RPE, its information could be easily used. Moreover, the integRPE algorithm requires value-decay for stability (Fig. 4Ba), and then, as discussed in the Results, value-decay could achieve simultaneous representation of the integrated TD-RPEs and the integrated state values. In situations where animals/humans need to quickly learn from a small number of samples and/or the environment is rapidly changing, accurate representation of the entire reward distribution may not be possible even by the segreRPE algorithm. Then, the simply implementable integRPE algorithm may suffice, as it can still learn reward distribution to a certain degree (Mikhael and Bogacz, 2016). The integRPE algorithm may thus be compatible with large learning rate and value-decay, as we suggested for CDh and its homolog DMS. This is potentially in line with the weaker distributional coding in DMS than in lAcbSh (Lowet et al., 2024).

Distributional coding was shown to be relatively weak also in DLS (Lowet et al., 2024). This apparently seems at odds with our suggestion of no value-decay in DLS, which is compatible with the segreRPE algorithm. However, the weak distributional coding in DLS can be due to incomplete learning because of small learning rate, as discussed above. Alternatively, it can be due to dominance of D2 over D1 values/RPEs, which was suggested for explaining the positive shift of RPE in DLS (Tsutsui-Kimura et al., 2020) and is potentially in line with the stronger projection from the primary motor cortex to D2 than D1 pathway (Wall et al., 2013; Lu et al., 2021).

## Acknowledgments

KM was supported by Grants-in-Aid for Scientific Research 23H03295, 23K27985, and 25H02594 from Japan Society for the Promotion of Science (JSPS) and the Naito Foundation. AK was supported by JSPS Overseas Research Fellowships.

## Extended Data legend

**Extended Data Figure 1-1.**
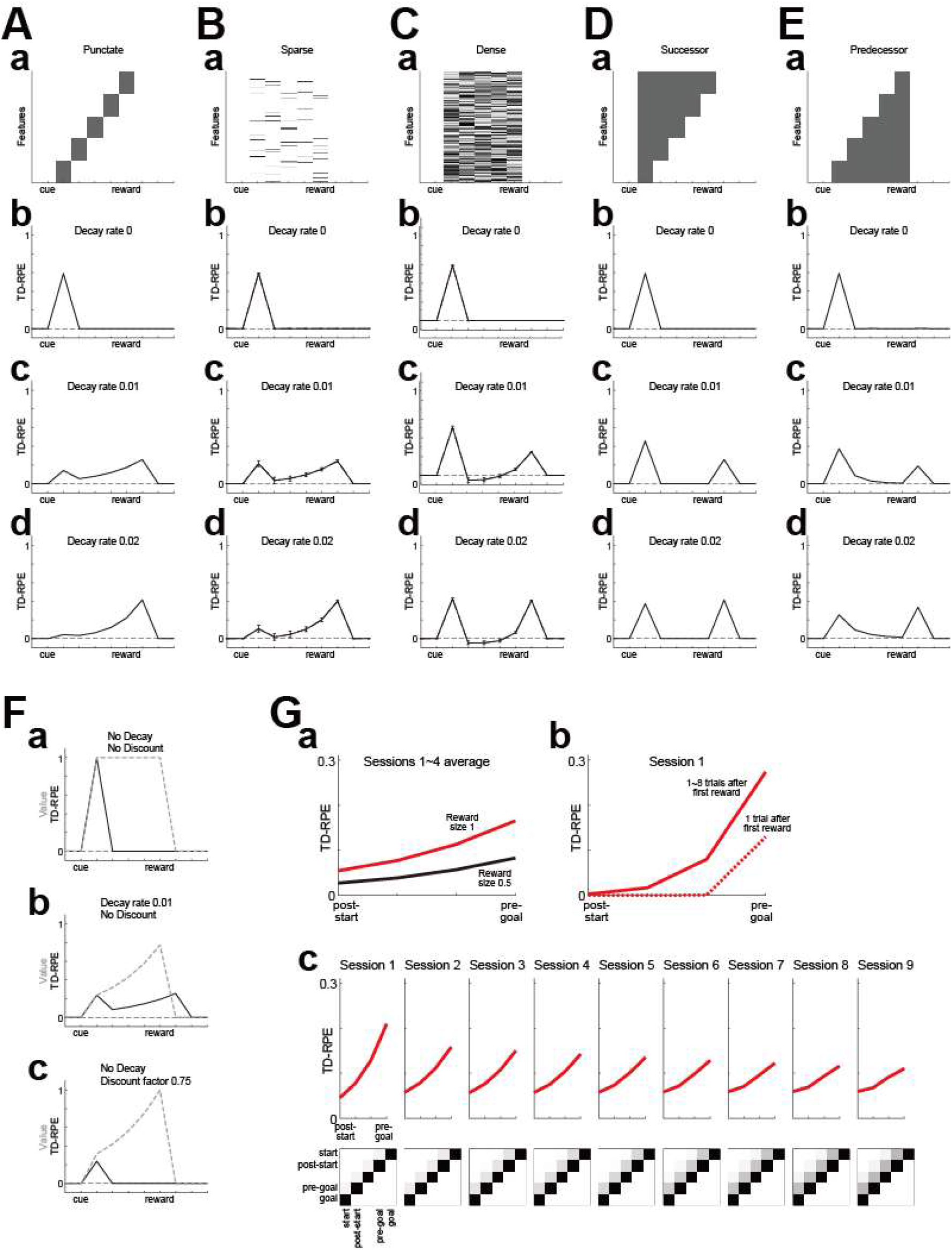
Results of simulations when the time discount factor *γ*, which was set to 1 in Figure 1 (except for those shown in Figure 1Fc), was instead set to 0.9.

**Extended Data Figure 1-2.**
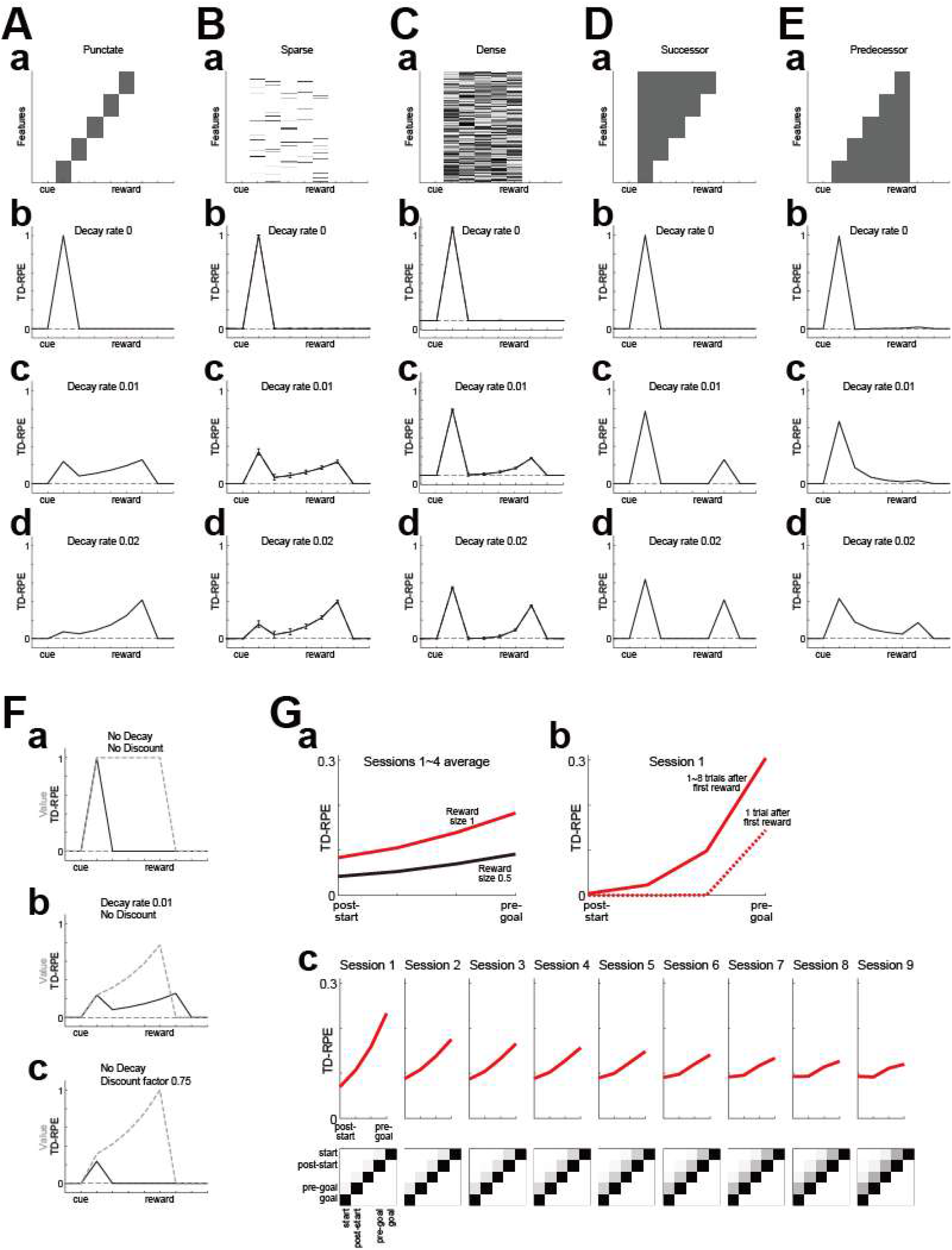
Results of simulations when (*a*_+_, *a*_−_), which were set to (0.15, 0.075) in Figure 1, were instead set to (0.15, 0.15).

**Extended Data Figure 3-1.**
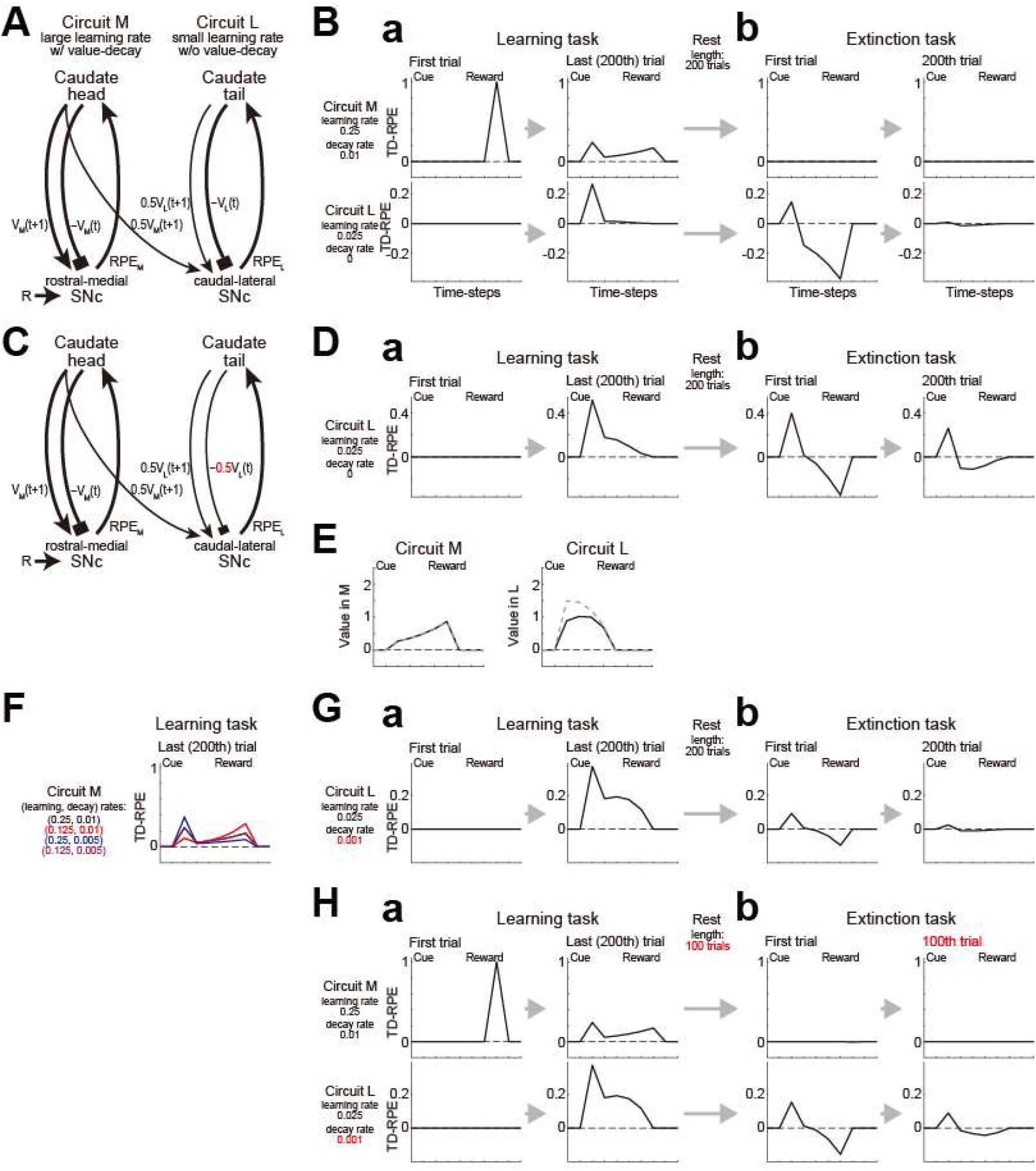
Results of simulations when the time discount factor *γ*, which was set to 1 in Figure 3, was instead set to 0.9.

**Extended Data Figure 4-1.**
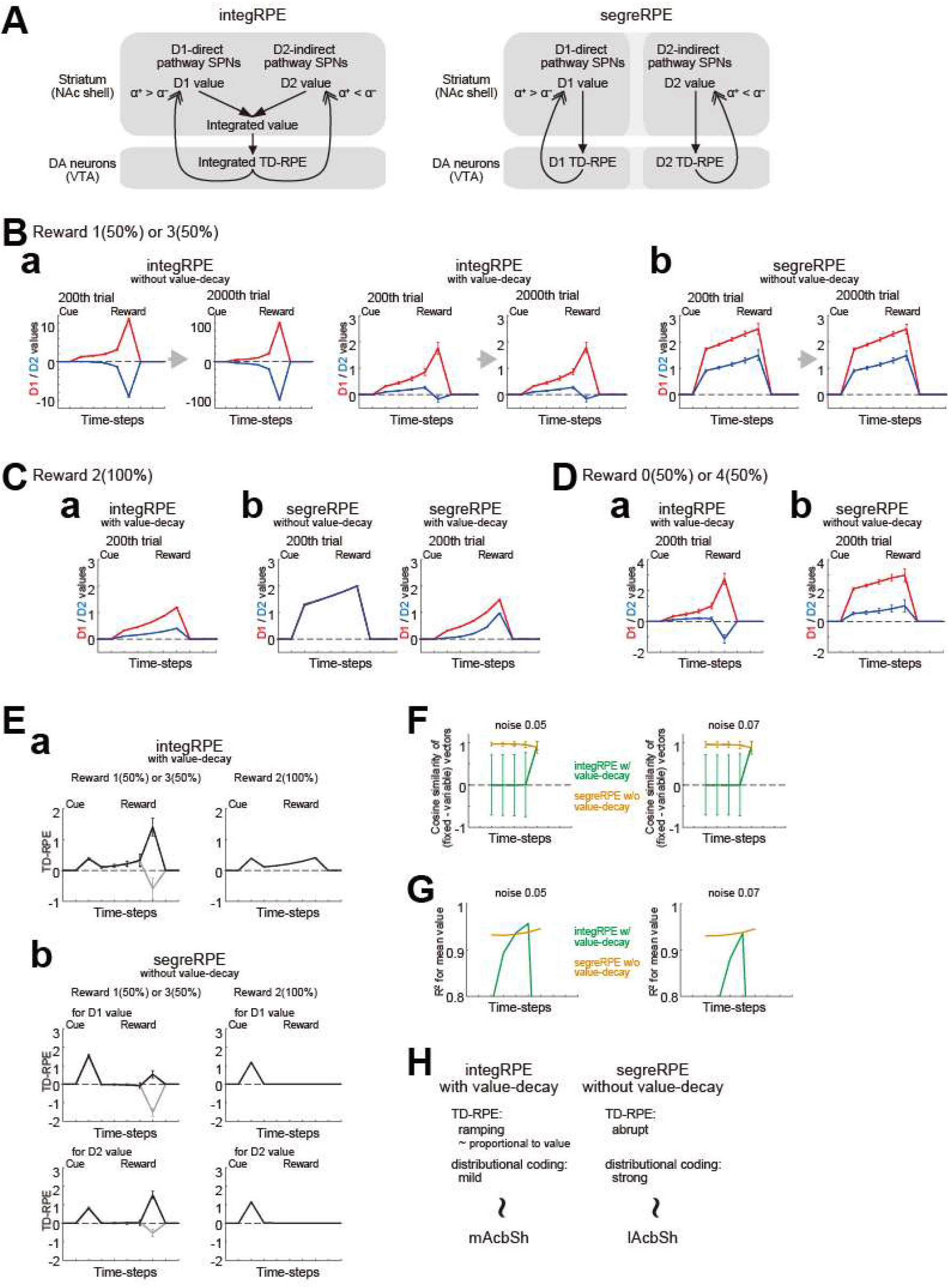
Results of simulations when the time discount factor *γ*, which was set to 1 in Figure 4, was instead set to 0.9.

**Extended Data Figure 5-1.**
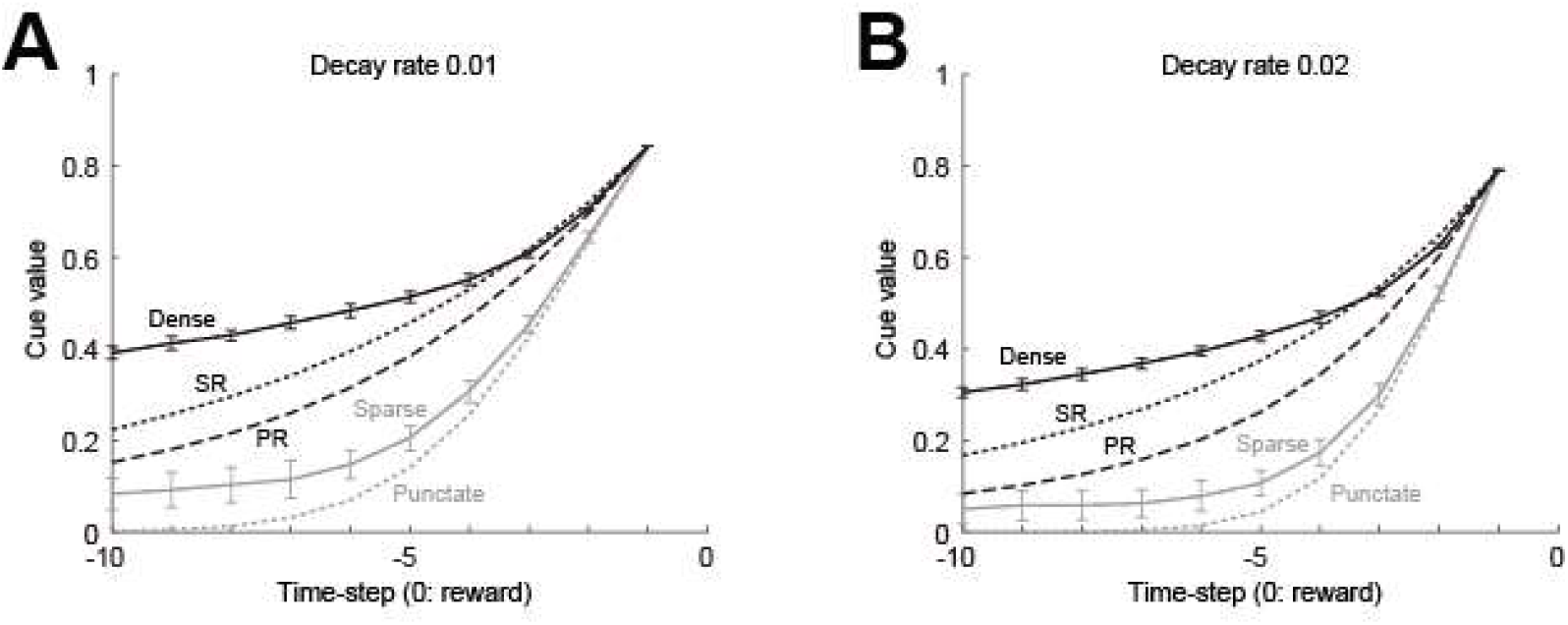
Results of simulations when the time discount factor *γ*, which was set to 1 in Figure 5, was instead set to 0.9.

**Extended Data Figure 5-2.**
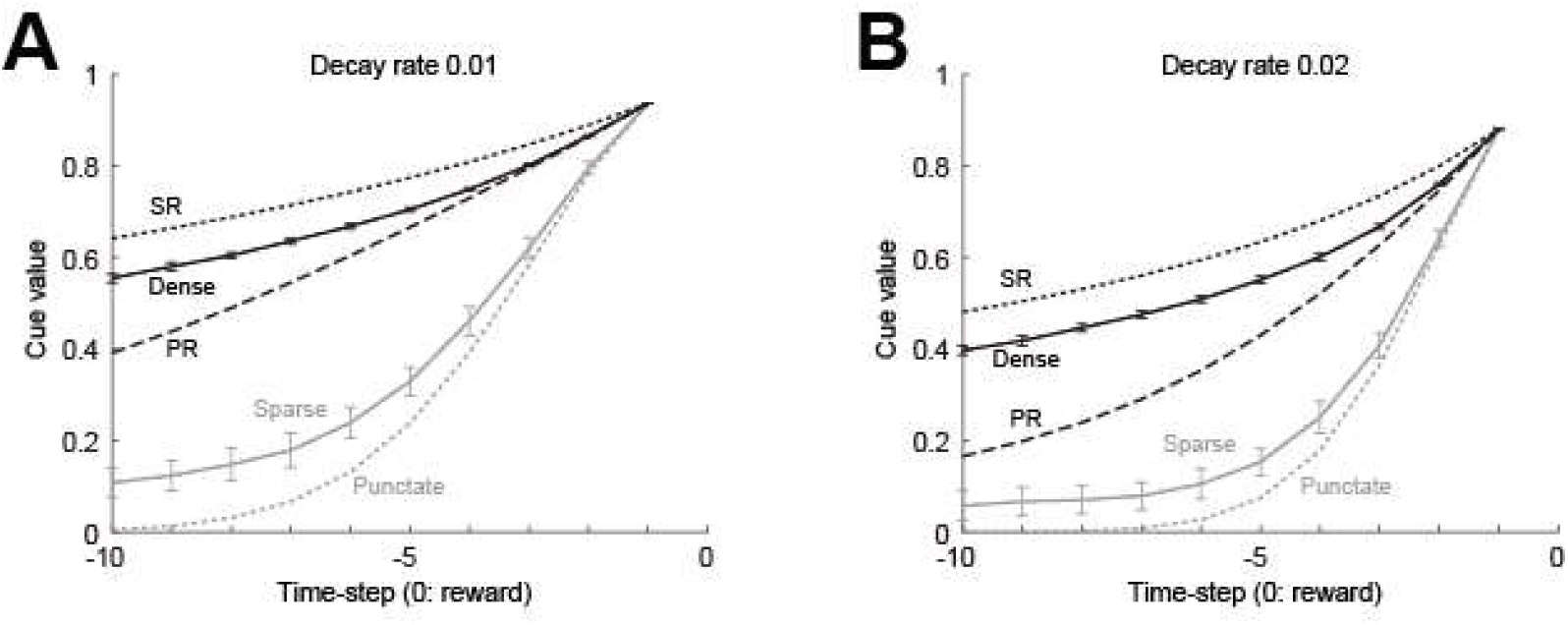
Results of simulations when (*a*_+_, *a*_−_), which were set to (0.15, 0.075) in Figure 5, were instead set to (0.15, 0.15).

**Extended Data Figure 6-1.**
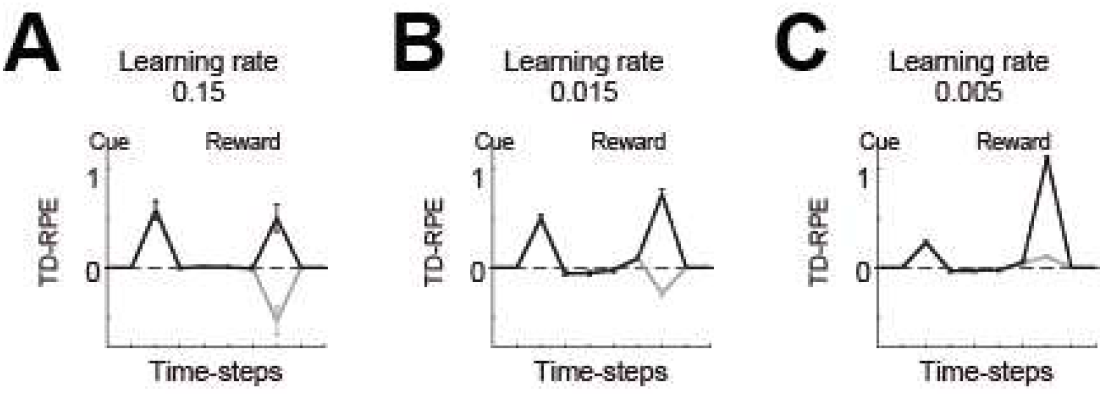
Results of simulations when the time discount factor *γ*, which was set to 1 in Figure 6, was instead set to 0.9.

